# Two types of regeneration mechanism in acute liver injury

**DOI:** 10.1101/2024.08.04.606468

**Authors:** Tomomi Aoyagi, Takeshi Goya, Koji Imoto, Yuki Azuma, Tomonobu Hioki, Motoyuki Kohjima, Masatake Tanaka, Yoshinao Oda, Yoshihiro Ogawa

**Author notes:** Corresponding author: Tanaka Masatake M.D., Ph.D. These two authors contributed equally to this study (Tomomi Aoyagi, Takeshi Goya).

## Abstract

The liver has a strong regenerative capacity, but the mechanisms of liver regeneration are not well understood. Furthermore, many previous studies on liver regeneration have been conducted in partial hepatectomy models, which may differ from acute liver injury with inflammation and necrosis, as observed in many clinical cases. In this study, we conducted a single-cell RNA-seq analysis (scRNA-seq) of liver regeneration in mice treated with acetaminophen (APAP) using publicly available data. We discovered that two cell proliferation populations appeared simultaneously during a single regenerative process. The two populations differed significantly in terms of differentiation, localization, proliferation rate, and signal response. Furthermore, one of the populations was induced by contact with necrotic tissue and exhibited a higher proliferative capacity with a dedifferentiated feature. These findings can shed new light on liver regeneration and aid in the development of therapeutic strategies for liver failure.

## Introduction

Liver disease (viral hepatitis, cirrhosis, and liver cancer) kills over 2 million people each year, accounting for 4% of global deaths ^1^. Therefore, a reduction in liver-related deaths is highly desirable. The liver has a high regenerative capacity and can return to its original size even after two-thirds of it is removed ^2^. However, the regeneration mechanism is not fully understood, and liver transplantation is the only treatment for liver failure. Therefore, clarification of the regeneration mechanism is urgently required.

Although the origin of regenerative cells in the liver is still debated, recent studies have suggested that the primary source of regenerative cells is preexisting mature hepatocytes rather than a small population of hepatic stem cells ^2 3^. Chen et al. found that the region where hepatocytes proliferate is determined by the type and extent of injury and that cells in all regions can proliferate ^4^. Furthermore, it has been reported that many genes associated with hepatocyte function are downregulated in proliferating hepatocytes ^5^. Chembazhi et al. found that hepatocytes in regeneration after hepatectomy appear to be divided into two types: proliferative hepatocytes and hypermetabolic hepatocytes that compensate for impaired liver function ^6^. Although previous studies with reporter mice revealed the localization of proliferation, the molecular biological mechanism in proliferating cells remains unknown. Furthermore, because liver injury is associated with hepatocyte destruction and inflammation, the regenerative mechanism of liver injury may differ from that of hepatectomy. However, a few studies have used scRNA-seq to investigate the mechanism of liver regeneration in the context of hepatocyte destruction and inflammation.

In this study, we conducted scRNA-seq of APAP-induced liver injury in mice using publicly available data and histological evaluation. These findings revealed the emergence of two proliferating cell populations during liver regeneration, with distinct differentiation states, localization, proliferation rates, and signal responsiveness. Notably, contact with necrotic tissue induced one of the proliferating cell populations, resulting in a more vigorous proliferative capacity and dedifferentiation. Furthermore, these cells were found in human acute liver injury. These findings may provide new insights into liver regeneration and aid in the development of future therapeutic strategies for liver failure.

## Results

### The specific cell population has high proliferative capacity in hepatic regeneration

To investigate transcriptomic changes in hepatocytes during regeneration, we performed scRNA-seq on APAP-treated mice using publicly available data ^7^. Clustering of the hepatocyte data yielded 14 clusters ranging from 0 to 13 (Fig. 1A). Clusters 7, 10, 12, and 13 were not found at 0 h and contained only a few cells that differed from the main population at each time point.

**Figure 1.**
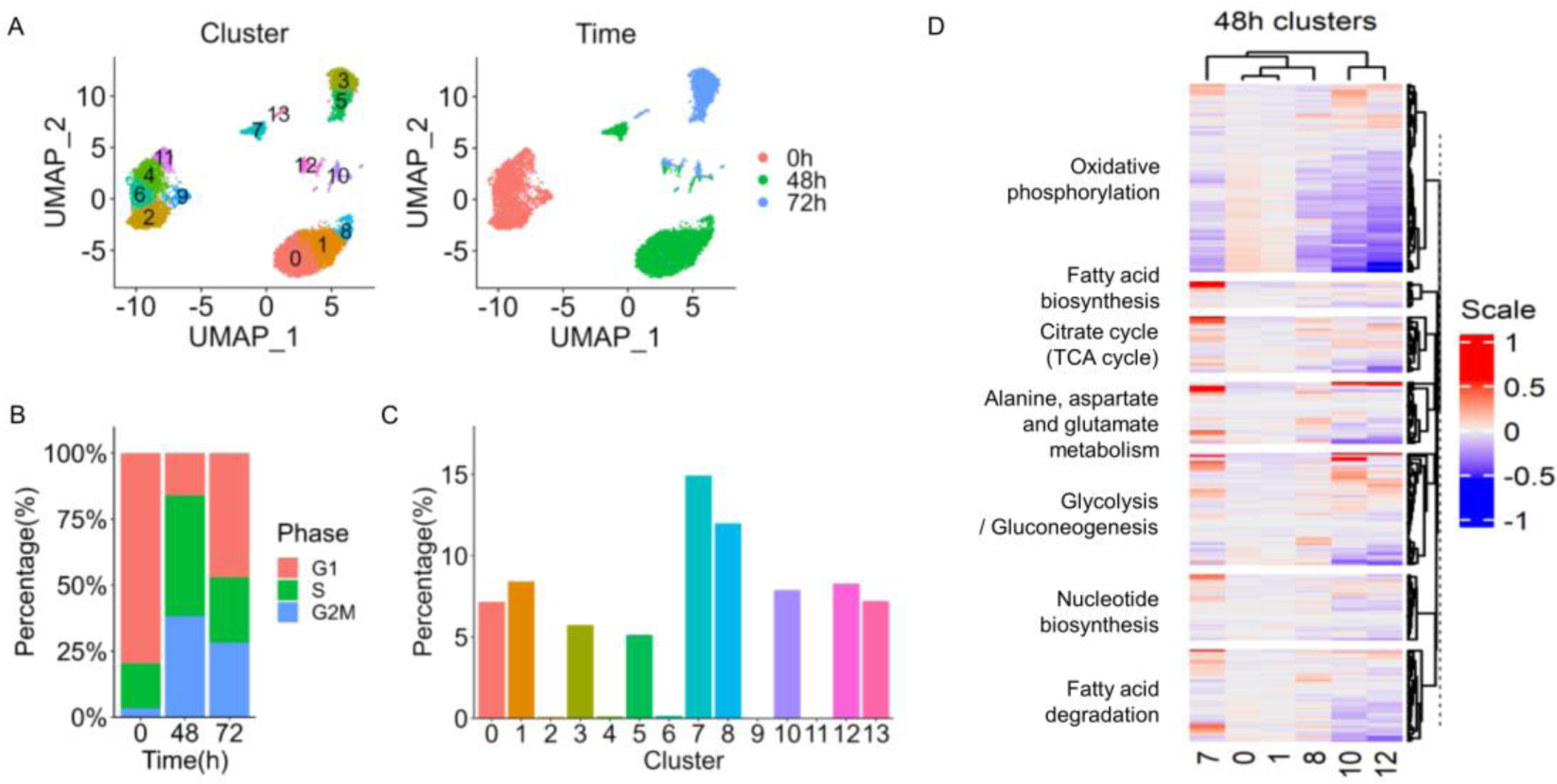
The specific cell population has high proliferative capacity in hepatic regeneration. (A) Uniform Manifold Approximation and Projection, with 10,114 hepatocytes integrated. Colors represent clusters and time after APAP administration. (B) The percentage of cells in their cycle phase at each time point. (C) The percentage of *Mki67*-positive cells in each cluster. (D) A heatmap showing the average expression levels of metabolism-related genes in 48-hour clusters.

The cell cycle analysis based on the gene expression profile revealed that the percentage of cells in the S and G2-M phases increased the most at 48 hours (Fig. 1B). The average expression levels of *Mki67*, *Pcna*, and *Tyms*, which are known to be upregulated in proliferating cells, increased after 48 hours (Fig.S1A). Therefore, we focused on clusters at 48 hours to investigate proliferating cells. To identify the cluster with the highest proliferative capacity, we compared the percentage of cells expressing *Mki67*, the gene for Ki67 (Fig. 1C). Cluster 7 had the highest percentage of *Mki67*-positive cells. Genes involved in nucleotide synthesis, glutamate synthesis, and energy metabolism, all of which are required for cell division, were upregulated highest in cluster 7 (Fig. 1D). Thus, cluster 7 was regarded as the most proliferative cell population.

### Two proliferating clusters emerge during the regeneration process

To better understand cluster 7, we further examined scRNA-seq data. Differentially expressed genes of cluster 7 were defined by the upregulation of genes involved in mRNA splicing such as *Malat1*, *Neat1,* and *Trp53inp1* (Fig. 2A, Fig. S2A). Based on the expression patterns of feature genes, clusters 7 and 13 were comparable populations, with cluster 7 being a 48-hour cluster and cluster 13 being a 72-hour cluster.

**Figure 2.**
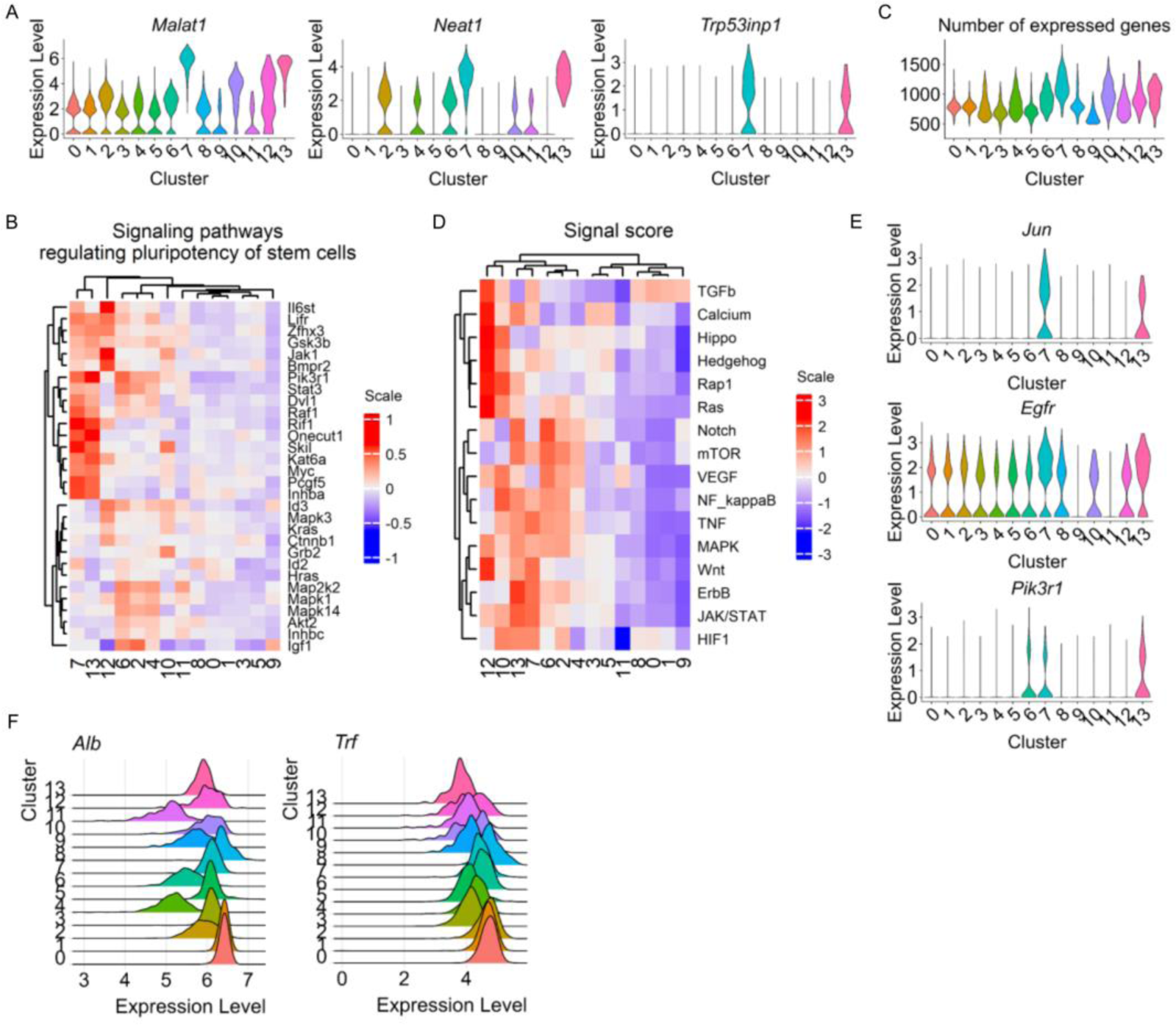
Two proliferating clusters emerge during the regeneration process. (A) Violin plot of the feature genes in cluster 7. (B) Heatmap of gene expression related to KEGG’s signal pathways that regulate pluripotency of stem cells. (C) A violin plot depicting the number of expressed genes in each cluster. (D) A heatmap depicting the average signal score for each cluster. (E) Violin plot of genes commonly expressed in the signal-enhanced cluster 7. (F) Ridge plots for *Alb* and *Trf* for each cluster.

Because proliferation is associated with dedifferentiation in many cell types, genes involved in stem cell regulation were investigated using the KEGG pathway database (mmu04550) (Fig. 2B) ^8 9,10^. In cluster 7, the expression of those genes was higher than in other clusters. Furthermore, Gulati et al. found that the number of expressed genes was higher in poorly differentiated cells, making it useful for predicting differentiation state ^11^. The number of expressed genes in each cluster was used to assess the differentiation level. The analysis revealed that cluster 7 had a greater number of expressed genes (Fig. 2C). These findings indicated that the cluster 7 cells exhibited characteristics of dedifferentiated cells.

To determine which signals are upregulated in cluster 7, we scored each signal using genes from the KEGG signal transduction pathway associated with cell proliferation. The findings revealed that signals such as ErbB and MAPK were upregulated in cluster 7 (Fig. 2D). Individual genes associated with these signals, including *Jun*, *Egfr*, and *Pik3r1,* were upregulated in cluster 7 (Fig. 2E).

Next, we investigated *Alb* and *Trf* expression to see if we could identify the hypermetabolic clusters described by Chembazhi et al. (Fig. 2F) ^6^. The results revealed that cluster 0/1/8, which contained the majority of 48 h cells, had higher *Alb* and *Trf* expression than the other clusters, including those in 0 h. However, many cells from cluster 0/1/8 expressed *Mki67* (Fig. 1C, Fig. S2B), indicating that they were also undergoing cell proliferation. We assessed and investigated the signals that regulated these phenotypes; however, the analysis of the scRNA-seq revealed no signals associated with cluster 0/1/8 (Fig. 2D).

These findings indicated that two proliferative cell populations emerged during the process of liver regeneration in the APAP model: one is represented by cluster 7, which has increased proliferative capacity due to dedifferentiation and increased *Egfr*/*Jun* expression. The other is cluster 0/1/8, in which cells proliferate as highly differentiated hepatocytes.

Cluster 10 expressed many genes with leukocyte characteristics, such as *Ptprc* and *Lyz2*, whereas cluster 12 expressed many genes associated with sinusoidal endothelial cells, such as *Clec4g* and *Pecam1*. Immunostaining revealed that no hepatocytes expressed the feature genes of clusters 10 and 12. Thus, we concluded that clusters 10 and 12 were contaminated with nonparenchymal cells.

### The cell population with high proliferation and dedifferentiation is the cells facing necrosis

For histological evaluation, mice were sacrificed at 12 h, 24 h, 48 h, and 72 h after APAP administration, and HE staining was conducted (Fig. 3A). Alanine aminotransferase (ALT) peaked at 24 h (Fig. S3A). In HE staining, necrosis appeared around the central vein at 12 h, followed by transient lipid accumulation in hepatocytes at 48 h, which decreased at 72 h (Fig. 3B). Hepatocytes with characteristic morphology emerged and localized in a layer of one cell-thickness bordering the necrotic area at 48 h. These peri necrotic cells had nuclei with prominent nucleoli and heterochromatin but did not accumulate lipids (Fig. 3B, arrows). After 72 h, these cells were still present in the necrotic area. To determine the location of cluster 7 cells, we conducted immunohistochemical staining of *Trp53inp1* and *Jun* proteins (TP53inp1 and c-jun), which were specifically expressed in cluster 7 in scRNA-seq (Fig. 3C, Fig. S3B, Fig. S3C). The results revealed that TP53inp1- or c-jun-positive cells were found facing the necrotic area, which corresponded to the characteristic peri necrotic cells observed by HE staining. Furthermore, the subcellular localization of TP53inp1 changed with time (Fig. S3C): cytoplasm at 12 h and nuclear membrane at 48 h. We named these hepatocytes with distinct morphology in the peri or intranecrotic layers as “Interfacing Necrotic area Transient Regenerative cells” (INTR cells) and investigated the positivity rate of cluster7 markers (TP53inp1 or c-jun) in the INTR cells (Fig. 3D). Approximately 80% or more of INTR cells expressed cluster 7 markers at all time points. These markers were expressed by almost no hepatocytes except INTR cells (data not shown). Therefore, we concluded that the cluster 7 cells were very similar to INTR cells. We also classified hepatocytes other than INTR cells as non-INTR cells, and because the 48 h clusters were divided into cluster 7 and cluster 0/1/8 in scRNA-seq, we concluded that cluster 0/1/8 was non-INTR cells. Following that, we identified INTR and non-INTR cells based on histological localization and cell morphology and classified them as clusters 7 and 0/1/8.

**Figure 3.**
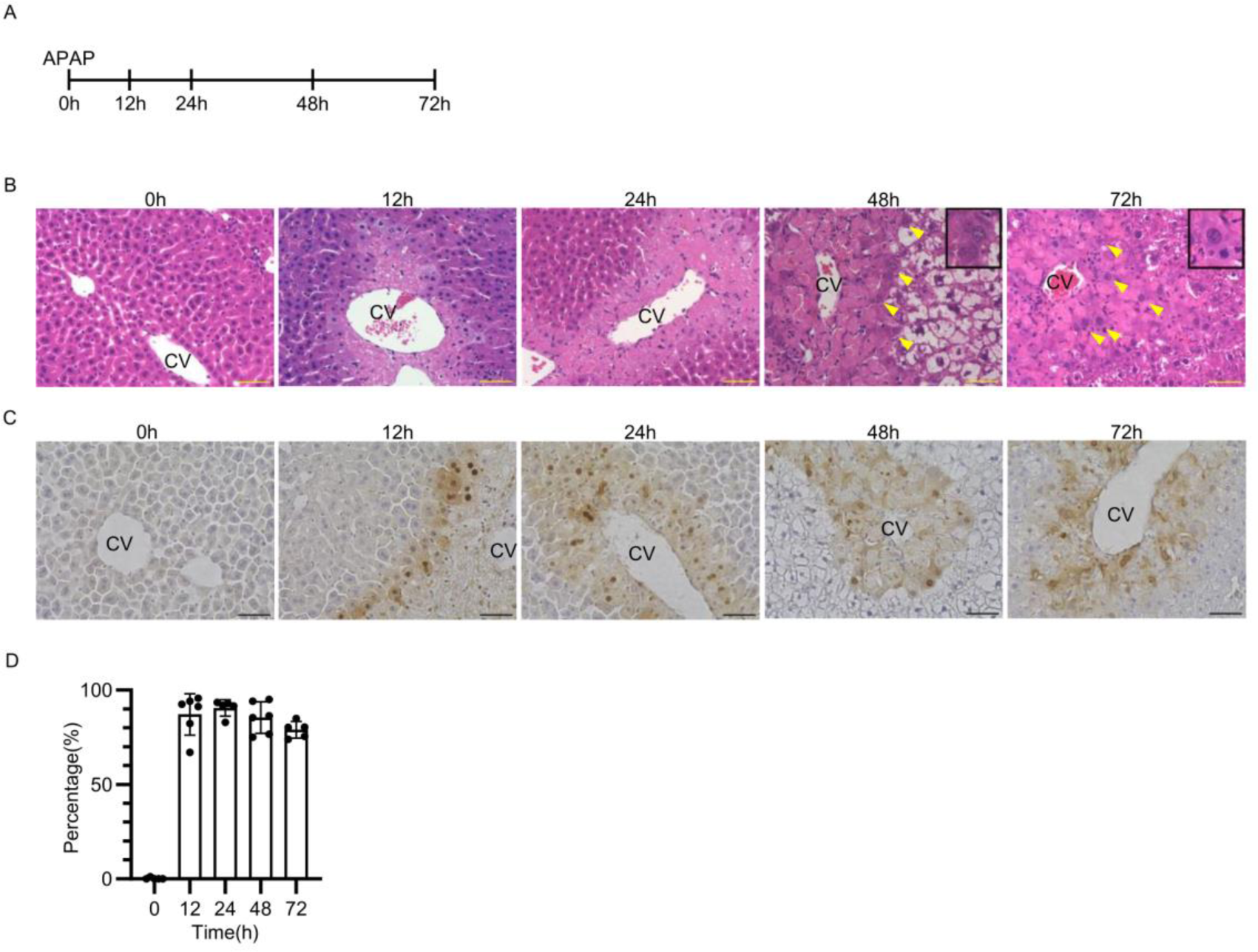
The cell population with high proliferation and dedifferentiation is the cells facing necrosis. (A) Shema for experimental design. Mice were sacrificed at the times specified. (B) HE staining at various time points. Arrows indicate cells with prominent nucleoli of interest. The scale bar is 50 µm. The inset shows representative cells of arrows. CV refers to the Central Vein. (C) Immunohistochemical staining of mixed antibodies for TP53inp1 and c-jun. The scale bar is 50 μm. CV, Central Vein. (D) Positive rate of TP53inp1 or c-jun in INTR cells.

Next, to see if INTR cells also appear in liver injury caused by other drugs, mice are given allyl alcohol (AA). Unlike APAP, administration of AA resulted in liver injury in the periportal region (Fig. S3D). Similar to APAP-induced liver injury, we discovered INTR cells bordering the necrotic area in AA-induced liver injury.

### INTR cells and non-INTR cells have different rates of cell cycle progression

To investigate changes in proliferative capacity over time, immunofluorescence staining for Ki67 was done (Fig. 4A, Fig. S4A). Consistent with the scRNA-seq results, both INTR and non-INTR cells contained Ki67-positive cells. INTR cells had a small number of Ki67-positive cells at 12 h and an increase in positive cells from 48 h to 72 h (Fig. 4B). The Ki67-positive rate among cells expressing TP53inp1, one of the markers of cluster 7, also increases at 72 h (Fig. S4B). In contrast, in non-INTR cells, Ki67-positive cells appeared after 48 h, with a lower positive rate than in INTR cells (Fig.4B). Thus, the time-dependent changes in Ki67-positive cells differed between INTR cells and non-INTR cells.

**Figure 4.**
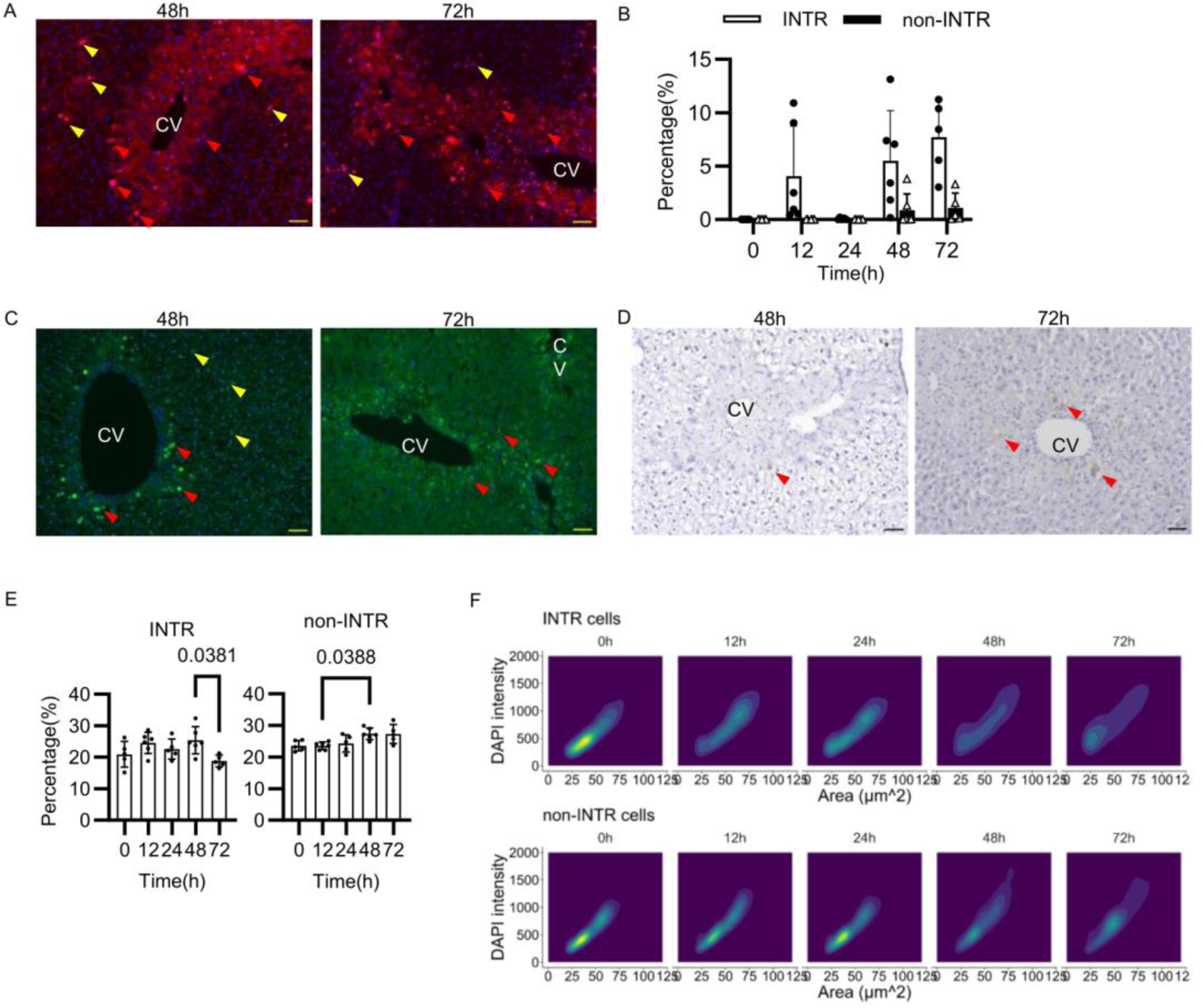
INTR cells and non-INTR cells have different rates of cell cycle progression. (A) Immunofluorescence staining for Ki67. The red arrowheads indicate representative positive cells of INTR cells. Yellow arrowheads denote representative positive non-INTR cells. TxRed, Ki67. Blue, DAPI. The scale bar is 50 μm. (B) Positive rate of Ki67. (C) Immunofluorescence staining for cyclinD1. The red arrowheads indicate representative positive cells of INTR cells. Yellow arrowheads denote representative positive non-INTR cells. The scale bar is 50 μm. (D) Immunohistochemical staining for cyclinB1. Red arrowheads denote representative positive cells. The scale bar is 50 μm. (E) The percentage of binuclear cells at each time point. (F) The distribution of nuclear size and DAPI intensity at each time point is shown. It was calculated using 2D kernel density estimation and depicted by a 2D density plot.

To determine the cell cycle status, immunostaining for cyclinD1 and cyclinB1 was used. CyclinD1 is known to appear in the G1/S phase of DNA replication, whereas cyclinB1 appears in the M phase of cytoplasmic division ^12^. CyclinD1 was strongly expressed in INTR cells after 48 h. In non-INTR cells, cyclinD1 was weakly expressed but in a greater number of cells than in Ki67 cells (Fig. 4C, Fig. S4C). On the other hand, cyclinB1-positive cells emerged in INTR cells from 48 h to 72 h, whereas no positive cells emerged in non-INTR cells until 72 h (Fig. 4D, Fig. S4D). These findings indicated that the rate of progression from the G1/S phase to the M phase varied between INTR and non-INTR cells. To determine the frequency of cytoplasmic division over time, we calculated the decrease in the number of binuclear cells, which corresponds to an increase in cytoplasmic division. We stained cell membranes with phalloidin and estimated the percentage of binuclear cells (Fig. 4E, Fig. S4E). In INTR cells, the rate of binuclear cells raised at 12 h, reduced at 24 h, raised again at 48 h, and fell again at 72 h. These findings showed that INTR cells divide for the first cytoplasmic division by 24 h, replicate DNA again at 48 h, and undergo a second cytoplasmic division by 72 h. The decrease of binuclear cells was greater from 48 h to 72 h than from 12 h to 24 h. Meanwhile, non-INTR cells increased their percentage of binuclear cells from 48 h to 72 h without decreasing. The cyclins and the percentage of binuclear cells show that INTR cells have two peaks of cytoplasmic division by 72 h, whereas non-INTR cells do not reach cytoplasmic division until 72 h.

Next, cell cycle progression was investigated using DAPI intensity and nuclear area. DAPI is a DNA stain, and its intensity and nuclear area are proportional to the amount of DNA, indicating cell cycle phase and polyploidy ^13 14^. Therefore, we estimated the amount of DNA using DAPI intensity and nuclear area to determine the difference in replication between INTR and non-INTR cells (Fig. 4F). Generally, because most cells are diploid and in the G0 and G1 phases, DAPI intensity and nuclear area cluster at a specific value. G2-M phase cells have twice as much DNA as G0/G1 phase cells, so they cluster twice as much. In the case of S-phase cells, the amount of DNA varies with S-phase progression, so they are sparsely distributed between the G1/G0 and G2-M populations. Because hepatocytes are polyploid in nature, some populations indicate tetraploidy or more, regardless of cell cycle. In both INTR and non-INTR cells, at 0 h, many cells cluster in an area of 30 μm^2^, indicating diploidy, while some cells, which appear to be tetraploid, cluster in an area of 60 μm^2^. In INTR cells, at 12 h, a cluster of cells shifts to 60 μm^2^, and at 24 h, the cluster splits into two areas of 30 μm^2^ and 60 μm^2^, indicating nuclear division. At 48 h, cell clustering disappears and cells are evenly distributed from 30 to 75 μm^2^, indicating an increase in the number of S-phase cells. At 72 h, the cell population increased again to 30 μm^2^, indicating a second nuclear division. This result indicates that INTR cells have two nuclear division peaks by 72 h, one at 12–24 h and one at 48–72 h, which is consistent with the percentage of binuclear cells. On the other hand, non-INTR cells remain nearly unchanged from 0 h until 24 h. At 48 h, the clusters start to fade, indicating an increase in S-phase cells. At 72 h, many cells increase their DNA content. In other words, non-INTR cells begin to replicate at 48 h, and many of them continue to replicate.

These findings show that INTR cells and non-INTR cells differ not only in localization but also in the rate of cell cycle progression, implying that INTR cells proliferate more quickly than non-INTR cells.

### Inhibition of EGFR, c-jun and mTOR significantly affect INTR cell proliferation

We assessed the role of *Egfr* and *Jun*, which were upregulated in cluster 7. The expression peak of *Jun* was earlier than that of *Egfr* in qPCR (Fig. S5A). To investigate the effect of EGFR, canertinib dihydrochloride, an irreversible inhibitor of EGFR phosphorylation, was administered. Bhushan. et al. reported that early inhibition of EGFR following APAP administration reduces liver damage, so canertinib dihydrochloride was injected 24 h after APAP administration, and mice were sacrificed 48 h later ^15^. The ratio of pEGFR to EGFR decreased in mice treated with canertinib (EGFRi group) (Fig. S5B, Fig. S5C). ALT levels did not differ between the EGFRi and control groups (Fig. S5D). In INTR cells, cyclinD1 positivity was not decreased, but Ki67 positivity was significantly reduced in the EGFRi group (Fig. 5A, Fig. S5E). In non-INTR cells, cyclinD1 and Ki67 positivity decreased, but not significantly. There was no difference in *Jun* expression between the EGFRi and control groups (Fig. 5B).

**Figure 5.**
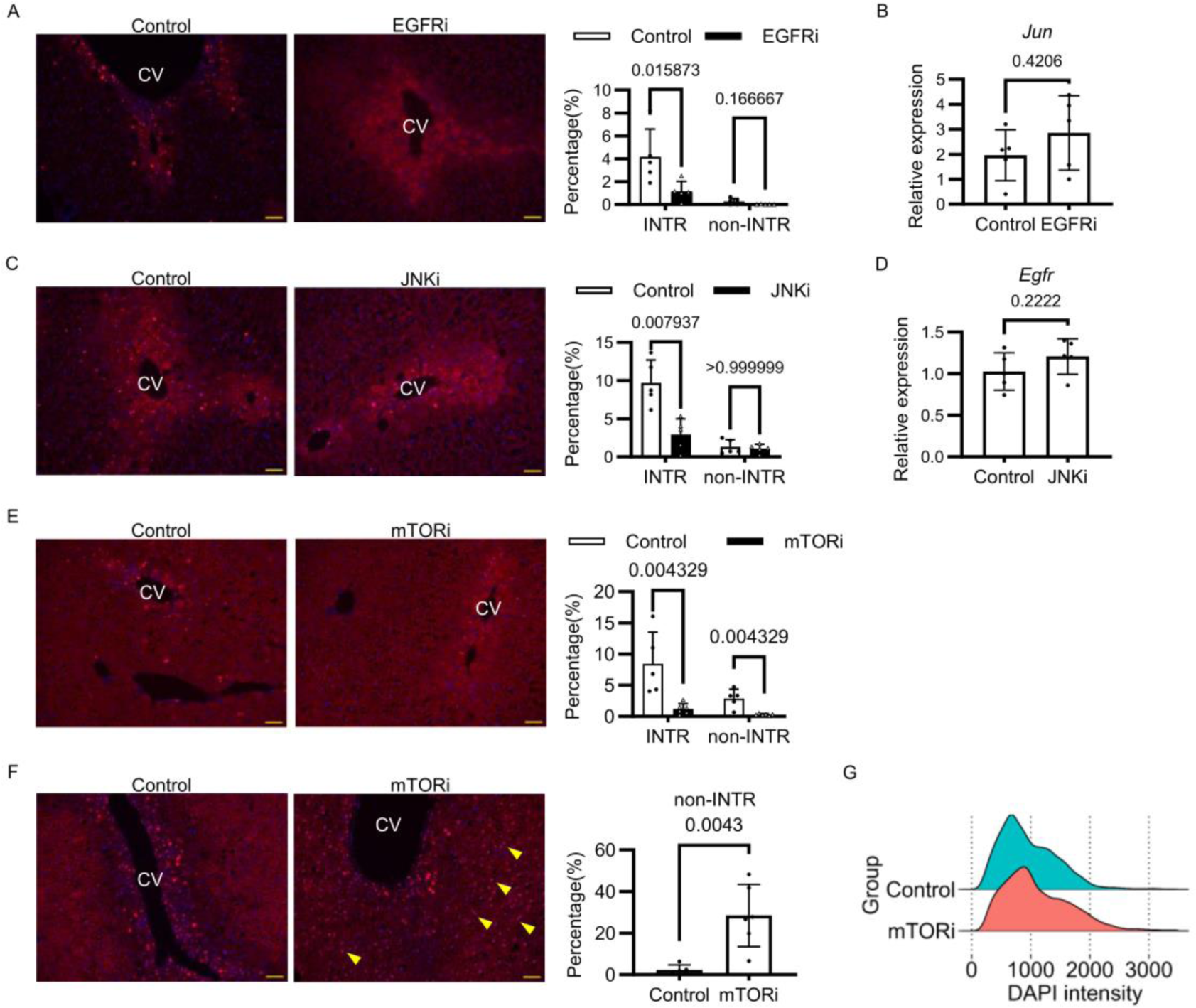
Inhibition of EGFR, c-jun and mTOR significantly affect INTR cell proliferation. (A) Immunofluorescence staining of Ki67 in an EGFR inhibition experiment. TxRed, Ki67. Blue, DAPI. The scale bar is 50μm. The bars represent the percentage of Ki67 positivity. (B) Relative mRNA expression of *Jun*, normalized by *Gapdh*. (C) Immunofluorescence staining of Ki67 in the JNK inhibition experiment. TxRed, Ki67. Blue, DAPI. The scale bar is 50μm. The bars represent the percentage of Ki67 positivity. (D) Relative mRNA expression of *Egfr*, normalized by *Gapdh*. (E) Immunofluorescence staining of Ki67 during the mTOR inhibition experiment. TxRed, Ki67. Blue, DAPI. The scale bar is 50 μm. The bars represent the percentage of Ki67 positivity. (F) Immunofluorescence staining for c-jun in the mTOR inhibition experiment. The arrowheads indicate representative positive cells in non-INTR cells. TxRed, c-jun. Blue, DAPI. The scale bar is 50 μm. Bars indicate the positivity rate of c-jun in non-INTR cells. (G) Ridge plot comparing DAPI intensity of non-INTR cell nuclei in the control and mTORi groups.

SP600125, an inhibitor of JNK, the kinase of c-jun, was then administered to investigate the effects of c-jun. Gunawan. et al. reported that early inhibition of JNK reduces APAP-induced liver injury, so SP600125 was given 12 h after APAP injection, and mice were sacrificed at 48 h ^16^. The ratio of phosphorylated c-jun to c-jun was reduced in the SP600125 group (JNKi group) (Fig. S5F, Fig. S5G). There was no difference in ALT between the JNKi and control groups (Fig. S5H). In INTR cells, cyclinD1 and Ki67 positivity was significantly reduced in the JNKi group. In non-INTR cells, cyclinD1 was reduced but not significantly, and Ki67 was not different between the JNKi and control groups (Fig. 5C, Fig. S5I). There was no difference in *Egfr* expression between the JNKi and control groups (Fig. 5D). These findings indicated that *Egfr* and *Jun* regulate the proliferation of both INTR and non-INTR cells, with INTR cells being more significantly impacted.

Next, we looked for the signal that controls the transduction of non-INTR cells. However, scRNA-seq revealed no signals regulated specifically in cluster 0/1/8. In homeostatic conditions, hepatocytes undergo both proliferation and physiological functions, such as protein synthesis and detoxification, like cluster 0/1/8. Therefore, we focused on mTOR, which has been found to promote cyclinD1 expression in homeostatic hepatocytes and hepatectomized mice^17 18 19^. We investigated its effects using rapamycin, a mTOR inhibitor. Algfeley. et al. reported that DMSO used to dissolve rapamycin reduces the hepatic damage caused by APAP, so rapamycin was administered 2 h after APAP administration and sacrificed at 48 h ^20^. There was no difference in ALT levels between the rapamycin-treated group (mTORi group) and control groups (Fig. S5J). CyclinD1 was significantly reduced in INTR cells after mTOR inhibition. In contrast, cyclinD1 was decreased in non-INTR cells, but the difference was not significant (Fig. S5K). Ki67 was significantly reduced in both INTR and non-INTR cells (Fig. 5E). Surprisingly, non-INTR cells in the mTORi group began to express c-jun (Fig. 5F). To investigate the effect of c-jun expression in non-INTR cells on cell cycle progression, we measured DAPI intensity (Fig. 5G). The results showed that the mTORi group had more cells with greater DAPI intensity than the control group. In other words, the mTORi group demonstrated a faster increase in DNA content than the control group. These findings demonstrated that mTOR signals influenced the proliferation of both INTR cells and non-INTR cells. Furthermore, inhibiting mTOR signals in non-INTR cells caused compensatory upregulation of c-jun, resulting in faster DNA replication.

### The contact with necrotic tissue is necessary for the conversion to INTR cells

Treatment with inhibitors of EGFR, JNK, and mTOR did not impact the appearance of INTR cells during HE staining (Fig. 6A). Because INTR cells consist of a single layer of cells bordering necrosis, contact with necrotic tissue may trigger of conversion of INTR cells. As a resulted, we investigated the effect of necrotic tissue on hepatocytes. We homogenized one mouse’s liver and injected it into the liver of another mouse. As a result, cells in contact with the necrotic material exhibited positive INTR cell markers, which did not appear after gel administration (Fig. 6B).

**Figure 6.**
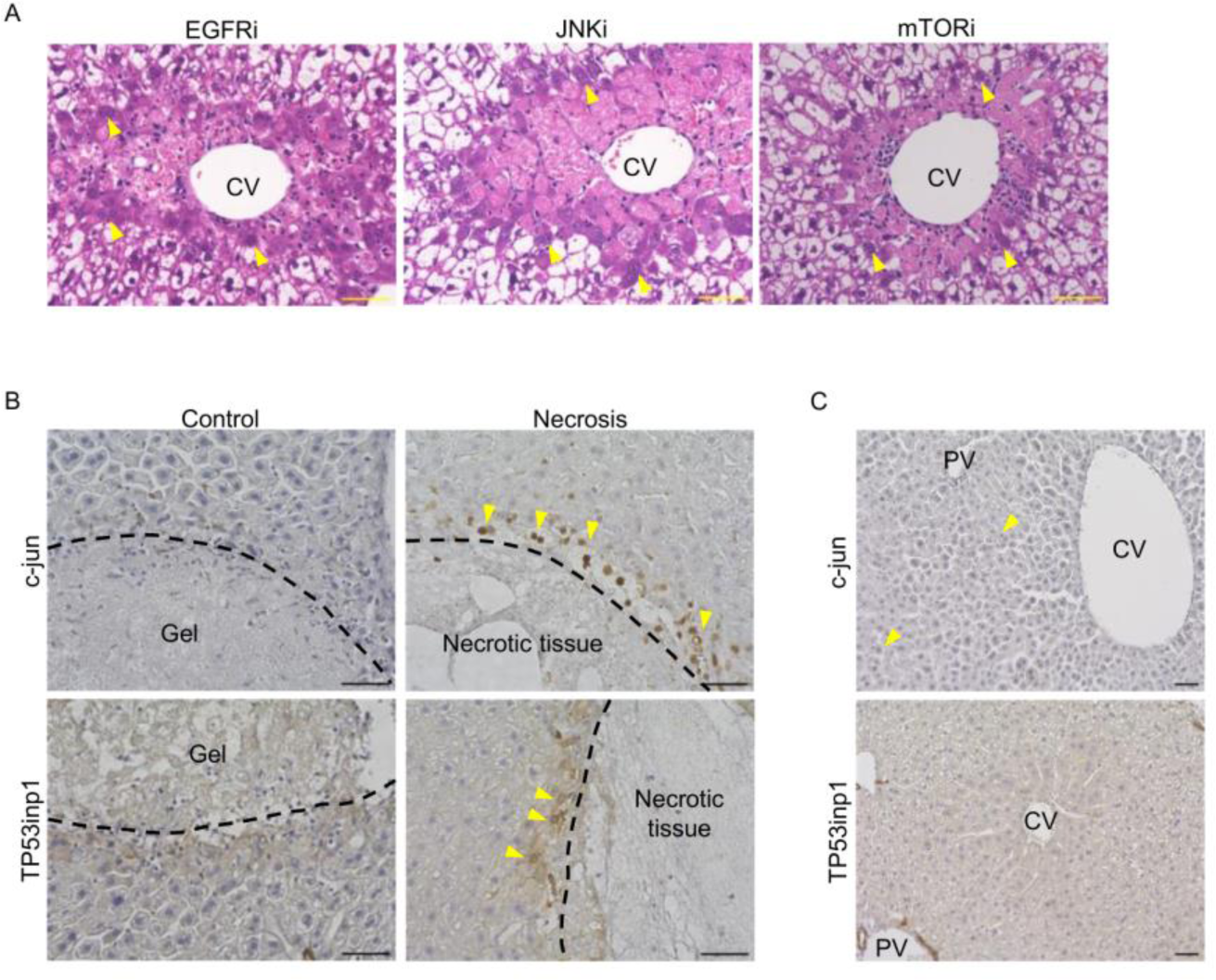
The contact with necrotic tissue is necessary for the conversion to INTR cells. (A) HE staining of mice treated with EGFRi, JNKi, or mTORi. The arrowheads indicate representative INTR cells. The scale bar is 50 μm. (B) Immunohistochemical staining for c-jun and TP53inp1 around necrotic tissue-injected or gel-injected areas. The arrowheads represent positive cells. The scale bar is 50 μm. (C) Immunohistochemical staining for c-jun and TP53inp1 in partially hepatectomized mice. The arrowheads represent positive cells. The scale bar is 50 μm.

Next, we examined the INTR cells in the hepatectomy model, which does not have inflammation or necrosis. We conducted a scRNA-seq analysis of the mice hepatectomy model using publicly available data ^6^. In this study, no cluster had an expression signature similar to INTR cells or upregulated INTR cell-specific genes like *Jun* and *Trp53inp1* (Fig. S6A, Fig. S6B, Fig. S6C).

Furthermore, immunohistological analysis confirmed the absence of TP53inp1-positive cells in the hepatectomy model. Although a few hepatocytes tested weakly positive for c-jun, the staining intensity was lower and more sporadic than in INTR cells in the APAP and AA models (Fig. 6C). We hypothesized that the weak expression of c-jun in the hepatectomy model is controlled by other mechanisms. Hence, these findings suggested that converting to INTR cells requires contact with necrotic tissue.

### INTR cells also appear in human liver injury

Finally, to see if INTR cells can be found in humans, we examined liver tissue from patients with acute liver injury due to hepatitis B virus (HBV), autoimmune hepatitis (AIH), and drug-induced liver injury (DILI) treated at our institution. c-jun or TP53inp1-positive cells were also seen around the necrosis (Fig. 7A). These findings demonstrated that INTR cells appear in human liver injury with necrosis, regardless of the cause.

**Figure 7.**
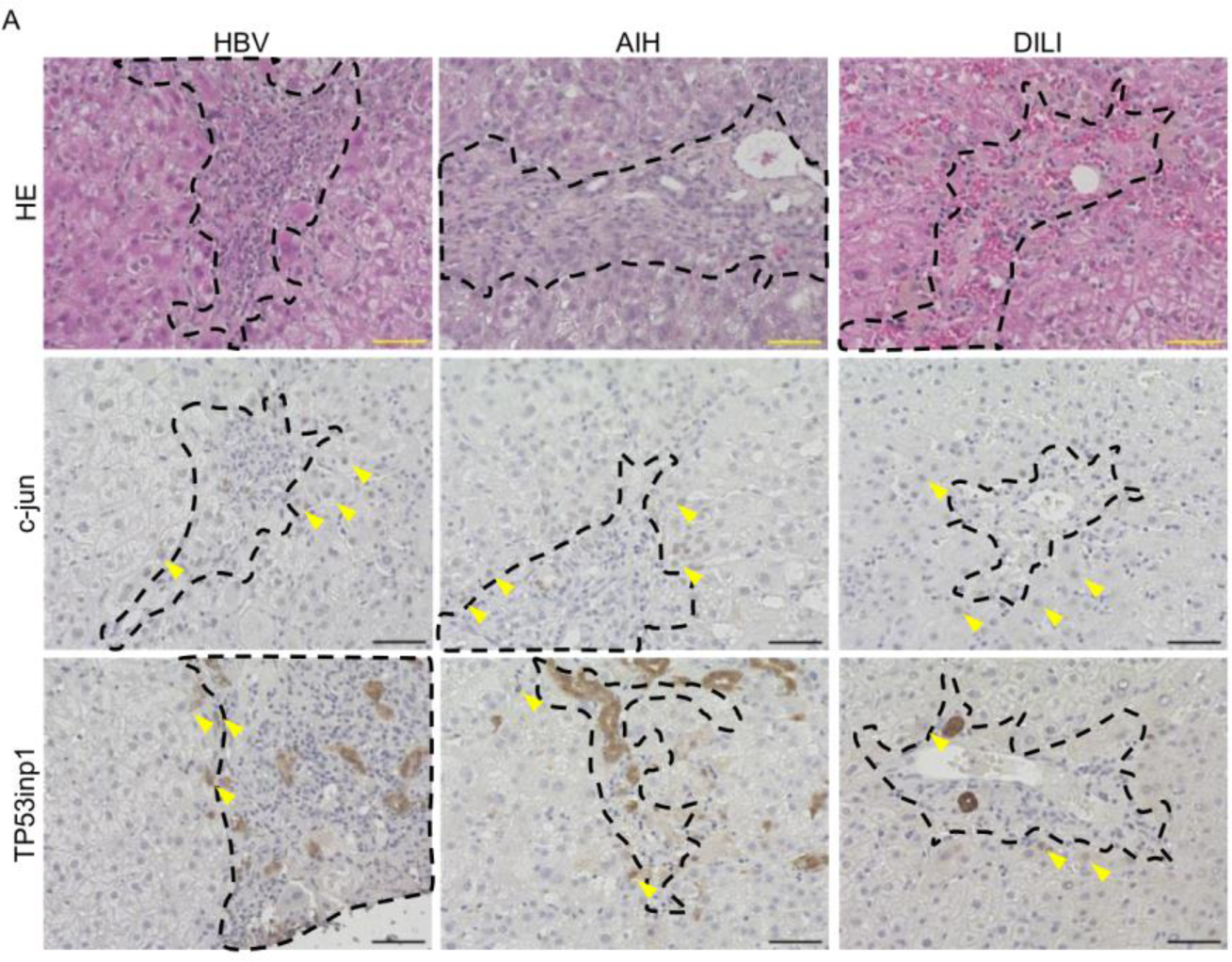
INTR cells also appear in human liver injury. (A) HE and immunohistochemical staining for c-jun and TP53inp1 in HBV, AIH, and DILI patients. The arrowheads represent positive cells. The inside of the dotted line represents the necrotic area. The scale bar is 50 μm.

## Discussion

In this study, we identified two distinct populations of proliferating cells that coexist during the regenerative process. It is known that cells with high proliferative capacity emerge during liver regeneration. Previously, many studies reported the various characteristics of proliferative hepatocytes, such as cell markers and signaling pathways^21 2^. However, these results were not consistent across the reports. Furthermore, although some of these cells in previous reports had characteristic localization depending on the type of liver injury, one layer of cells bordering necrotic tissue similar to INTR cells has not been reported. Recently, a study by Chen. et al., using rainbow mice, demonstrated that cell division occurs in all zones, not just one ^4^. Similarly, Wei. et al. and He. et al. reported the proliferative capacity in the entire lobule with regional differences^17,22^. Ben-Moshe. et al. also found Ki67-positive cells not only in the peri necrotic area but also in zones far from the necrotic site, indicating that hepatocytes in all zones are involved in regeneration ^7^. Thus, recent studies demonstrated that hepatocytes throughout the liver lobules contribute liver regeneration.

However, the heterogeneity of these proliferative cells had not been clarified. Here, we revealed the two distinct proliferative hepatocyte populations with different proliferative capacities during liver regeneration. Consistent with previous findings, we discovered that cell proliferation occurs throughout the liver in the APAP mouse model. Furthermore, we discovered that proliferating cells are divided into two distinct populations, each with varying levels of differentiation, localization, cell cycle progression, and signal response. We also showed that contact with necrotic tissue causes one of the proliferating cells to become more proliferative capacity than another population.

We discovered characteristic cell clusters associated with dedifferentiation during the regenerative process. It is well understood that tissue-resident stem cells are the primary cell source in rapidly self-renewing organs like the intestine, skin, and blood ^8 9 10^. Most tissue stem cells are classified as a dedifferentiated state because of the expression of stem cell markers and their high proliferative capacity. INTR cells also express stem cell markers and have a high proliferative capacity, similar to other tissue stem cells. However, INTR cells were a distinct cluster in the regenerative process and did not populate in homeostatic conditions. The liver may have a distinct regenerative system separate from tissue-resident stem cells. Hepatocytes are known to be highly plastic; INTR cells could be one of the transdifferentiated cells derived from mature hepatocytes on demand ^23^.

In addition, INTR cells increased the expression of *Egfr* and *Jun*. Ligands for EGFR are critical cytokines in liver regeneration. Although many studies have looked into the effects of cytokines on liver regeneration, most of them are auxiliary mitogenic factors, and inhibiting them causes regeneration to be delayed but eventually completed. In contrast, simultaneous inhibition of HGF and EGFR ligands can completely inhibit liver regeneration in hepatectomy mice, making it a "complete mitogen" ^24^. In particular, EGFR has a significant effect on DILI, such as APAP and paracetamol, and inhibiting EGFR alone almost eliminates liver regeneration ^24 25^. Consistently, inhibition of EGFR reduced cell proliferation of INTR cells (Fig. 5A, Fig. S5E). Furthermore, c-jun is a member of the proteins of the activator protein 1 (AP-1) complex, which promotes cell proliferation. It has been reported that EGFR promotes the expression of c-jun, while c-jun can also promote the expression of EGFR; however, inhibition of EGFR or JNK had no effect on the expression of another in our analysis (Fig. 5B, Fig. 5D) ^26 27^. These findings suggested that the EGFR or c-jun signal independently controlled the proliferation of INTR cells. Furthermore, while both signals were required for INTR cell proliferation, inhibiting them did not affect the emergence of INTR cells. Thus, these signals are simply “growth factors,” not the causal factor that causes cell conversion to INTR cells.

On the other hand, the other proliferative cluster, cluster0/1/8, is a population of cells that are highly differentiated (Fig. 2F). It is possible that the simultaneous entry of all hepatocytes into the proliferative phase causes temporary liver failure because many hepatocyte differentiation genes are downregulated in proliferating cells ^5^. Similarly, Chembazhi. et al. found that during regeneration after hepatectomy in mice, hepatocytes are divided into a proliferation-specific population and a metabolically active population that compensates for reduced liver function ^6^.

Cluster0/1/8 has higher levels of *Alb* and *Trf* than other clusters at all time points, indicating that cluster0/1/8 may be the metabolically activated population (Fig. 2F). Notably, unlike the hepatectomy model, cluster0/1/8 also expressed numerous genes involved in cell proliferation (Fig. 1C, Fig. S2B). Regeneration after hepatectomy was not accompanied by hepatocyte destruction or inflammation, which could explain the difference from the hepatectomy model. Generally, conflicts between DNA polymerase and RNA polymerase caused by simultaneous replication and transcription increase genomic instability, such as DNA breaks ^28^. Therefore, it is reasonable for proliferating cells to dedifferentiate to reduce the risk of genomic instability. However, cluster0/1/8 developed both highly differentiated activity and cell proliferation capacity. These cells’ emergence may be an abnormal and unavoidable response to “emergencies.” In clinical settings, acute liver injury is frequently associated with inflammation and hepatocellular destruction, implying that our findings are more representative of clinically acute liver injury. Furthermore, the simultaneous acquisition of differentiation and cell proliferation is likely a temporary emergency response that ends with liver regeneration. Chronic inflammation may contribute to the persistence of this abnormal state. Thus, these non-INTR cells may be associated with genomic mutations and carcinogenesis during chronic liver injury, providing a key to understanding the mechanism of liver carcinogenesis. We need to conduct more research into chronic liver injury.

The two cell populations were clearly separated by their localization. Interestingly, the morphology of these two cell populations varied significantly. This finding is consistent with that of scRNA-seq, which suggested that the two populations had distinct characteristics because they were widely separated in Uniform Manifold Approximation and Projection. Ben-Moshe. et al. described INTR cells as "interfacing hepatocytes" ^7^. The "interfacing hepatocyte" exhibits increased expression of fetal markers, which is consistent with our findings in INTR cells. They reported that "interfacing hepatocytes" are pushed into the perivascular zone by surrounding cell proliferation and become less differentiated, transforming into perivascular zone-specific cells. Our findings suggest that interfacing hepatocytes are not only transformed into zone characteristics but also actively participate in hepatocyte proliferation. Furthermore, the 72 h HE staining suggested that INTR cells were actively infiltrating the necrotic area rather than being pushed forward (Fig. 3B).

TP53inp1, which is used as a marker for cluster 7, has been identified as an autophagy protein. TP53inp1 is a downstream protein of p53 that, when localized in the nucleus, exhibits apoptotic functions. However, when localized in the cytoplasm, it binds to LC3 and regulates autophagy. It is particularly associated with mitophagy and mitochondrial quality control ^29^. In our study, we also found TP53inp1 sporadically in the cytoplasm of INTR cells after 12 h, indicating mitophagy (Fig. S3C). This finding is consistent with previous reports that mitophagy occurs in peri necrotic cells during APAP liver injury ^30 31^. Consistent with reports that TP53inp1 is degraded during autophagy progression, TP53inp1 expression decreases at 24 h (Fig. S3C) ^32^. However, it is re-expressed at the nuclear membrane at 48 h and localizes to the cytoplasm at 72 h (Fig. S3C).

Given its nuclear membrane localization, TP53inp1 at 48 h is unlikely to function as a mitophagy. Unlike general autophagy, nuclear membrane autophagy is induced by Ras gene activation rather than starvation, and it results in cellular senescence as a tumor suppressor mechanism ^33^. Moreover, Umbaugh, D. S. et al. reported that expression of p21, a marker of cellular senescence, is increased in cells surrounding APAP-induced liver necrosis ^34^. INTR cells are perinecrotic and have upregulated EGFR expression. Because EGFR activates Ras, TP53inp1 at 48 hours may be involved in cellular senescence for tumor suppressor mechanisms via nuclear membrane autophagy. Thus, TP53inp1 in INTR cells may be involved in two types of autophagy: mitophagy in the cytoplasm at 12 h and nuclear membrane autophagy at 48 h.

INTR and non-INTR cells progress through the cell cycle at different rates. Although the presence of proliferating cells in liver regeneration has been demonstrated in previous studies using proliferation markers or transgenic mice, there has been no evidence that multiple distinct cell types with different cell cycle progression rates contribute to liver regeneration during the simultaneous regeneration process. It is difficult to accurately assess the rate of cell cycle progression, but we estimated it using four criteria: Ki67 positivity, cyclin expression pattern, number of nuclei, and DAPI intensity. In all experiments, INTR cells showed two mitotic peaks up to 72 h after observation, indicating a rapid progression of the cell cycle (Fig.4E, Fig.4F). In contrast, non-INTR cells began to enter S phase after 48 h (Fig.4F). In this study, it was unclear why non-INTR cells had a slower cell cycle rate, but when they switched to the c-jun pathway in the mTOR inhibitor experiment, they progressed faster than the control group. This finding implies that non-INTR cells can accelerate the cell cycle but are hesitant to do so. The slower rate of cell cycle progression may benefit non-INTR cell proliferation and hepatocyte function.

We identified three signaling pathways (EGFR, JNK, and mTOR) that regulate proliferation in INTR or non-INTR cells; however, these pathways are not involved in cell conversion and do not completely control proliferation. The inhibition of each signal affected both INTR cells and non-INTR cells. However, INTR cells and non-INTR cells responded differently to inhibitors. When one signal was inhibited, INTR cells produced significantly fewer Ki67- and cyclinD1-positive cells.

This finding suggests that each signal is critical to the proliferation of INTR cells. On the other hand, non-INTR cells showed a trend of decreasing cyclinD1-positive cells, but this decrease was not significant with any inhibitor. In addition, c-jun, which is normally expressed only in INTR cells, was also expressed in non-INTR cells after treatment with a mTOR inhibitor. In other words, non-INTR cells typically use the mTOR pathway, but when that pathway is disrupted, they quickly switch to the c-jun pathway. Unlike INTR cells, non-INTR cells can easily compensate for signal blockage with another signal.

We discovered that contact with necrotic tissue caused hepatocytes to convert into highly proliferative cells known as INTR cells. Inhibition of EGFR, JNK, and mTOR decreased the positivity of proliferation markers but did not reduce the number of INTR cells on HE staining.

EGFR, JNK, and mTOR are involved in INTR cell proliferation but do not serve as initiation signals. If the signal that triggers the change to INTR cells is a result of paracrine action, then 2–3 layers of cells from necrotic tissue can become INTR cells, depending on the signal strength. However, INTR cells are composed of a single layer of cells in contact with necrosis. Therefore, we hypothesized that contact with necrotic tissue is essential for initiating the transformation to INTR cells. Injection experiments with necrotic tissue revealed that it is necessary for the induction of INTR cells.

Furthermore, cluster 7 marker-positive cells were observed only found in cells that came into contact with necrotic tissue, indicating that this contact is critical. Furthermore, because necrotic tissue is injected percutaneously, it is randomly distributed. This result implies that all cells, not just those in a specific area, can become INTR cells.

Finally, it is possible that these regenerative mechanisms function in a wide range of liver injuries and are conserved across species. The focus of this study was on APAP-induced liver injury. However, INTR cells were also found in liver injury caused by AA, which has a different injury site than APAP. This finding suggests that the regenerative mechanism in our study is effective regardless of the cause of liver injury. Furthermore, c-jun or TP53inp1-positive cells were found in human acute liver injury caused by HBV, autoimmune hepatitis, and drugs. Because the exact onset time and necrosis boundaries are not visible in human liver injury, not all cells bordering the peri necrotic area were positive for the INTR cell marker, unlike in mice. However, as in mice, positive cells appeared primarily in the peri necrotic area, indicating that the regenerative mechanism described in this study is also active in humans.

Although we discovered the aforementioned liver regeneration mechanisms, we cannot identify the master regulator of INTR and non-INTR cells. We demonstrated that necrotic tissues induced INTR cells; however, it is unknown which molecule induced INTR cells. Furthermore, we identified several signaling pathways involved in proliferation; however, inhibition of these signals did not completely prevent liver regeneration. Probably, these signals are simply downstream effectors of the regulatory network. To develop novel therapeutic strategies for acute liver injury, we must identify more upstream factors that influence transcriptional regulatory networks. For more precise investigations, genes capable of lineage tracing should be defined; however, we cannot specify such genes. Moreover, it is reported that hepatic nonparenchymal cells, such as Kupffer cells and stellate cells, were involved in liver regeneration ^35,36^. Our analysis could not detect this cell-cell crosstalk, however it is possible that hepatic nonparenchymal cells influence the conversion to INTR or non-INTR cells and proliferation. Further research focusing on interaction between INTR or non-INTR cells and hepatic nonparenchymal cells are needed.

Nonetheless, we discovered a heterogeneous proliferative system in liver regeneration and necrotic tissue induced cell conversion with high proliferation potential. These findings may provide new insights for basic science research and the development of novel therapeutic strategies for liver failure.

## Acknowledgments

We would like to thank Makiko Kamihashi for their excellent technical assistance. We would like to thank Enago (www.enago.jpÄb0) for the English language review.

## Author contributions

Author Contributions: T.A., T.G., and M.T. designed the research studies. T.A. and T.G. performed experiments. T.A. and Y.O. (Yoshinao Oda) collected and analyzed data. T.A. and T.G. authored the manuscript. K.I., Y.A., T.H., and M.K. assisted with data analyses. M.K., M.T., and Y.O. (Yoshihiro Ogawa) helped analyze and interpret the data. T.G., M.T., and Y.O. (Yoshihiro Ogawa) helped prepare the manuscript and critically reviewed it. All authors have read and approved the published version of the manuscript.

## STAR Methods

### Animal models

Male C57B/6 J mice, 8 weeks old, were obtained from Japan SLC, Inc. (Shizuoka, Japan) and housed in a controlled environment with free access to food and water. They were kept in a standard cage with a 12-hour light/dark cycle, a normal room temperature of 25°C, and a relative humidity of around 50%. Before each experiment, the mice were randomly assigned to groups (n = 5–6 per group) and fasted overnight. In the APAP experiments, APAP (300 mg/kg) (Sigma-Aldrich Japan G.K., Tokyo, Japan; # A7085) was dissolved in warm phosphate-buffered saline (PBS) and administered intraperitoneally (i.p.) in the morning. For time series experiments, mice were sacrificed at 0 h, 12 h, 24 h, 48 h, and 72 h after APAP. For EGFR inhibition experiments, canertinib dihydrochloride (80 mg/kg, i.p.) (Selleck Chemicals, Tokyo, Japan; # S0711) was dissolved in PBS.

The mice were given canertinib dihydrochloride or PBS 24h after APAP, and they were sacrificed 48 h later. To conduct JNK inhibitor experiments, SP600125 (10 mg/kg, i.p.) (Selleck Chemicals, Tokyo, Japan; # S1460) was dissolved in 5%DMSO. SP600125 or 5% DMSO was administered 12 h after APAP, and mice were sacrificed 48 h later. For mTOR inhibitor experiments, rapamycin (2 mg/kg, i.p.) (Selleck Chemicals, Tokyo, Japan; # S1039) was dissolved in 2% DMSO. Rapamycin or 2% DMSO was administered 2 h after APAP, and the mice were sacrificed 48 h later. For necrotic tissue injection experiments, roughly 500 μl of liver from another mouse was incubated at 37°C for 2 hours and homogenized with 115 μl of PBS and 10 μl of sumi ink (as about 75% of the total consists of liver tissue). To make it gelatinous, carboxymethyl cellulose sodium (Sigma-Aldrich Japan G.K., Tokyo, Japan; # 05-1760) was incorporated at 1%–5% depending on the hardness. The prepared necrotic tissue or 5% carboxymethyl cellulose sodium (dissolved in PBS) was injected percutaneously into the mouse liver and sacrificed 48 h after later. In the AA administration experiment, 1% AA (FUJIFILM Wako Pure Chemical Corp., Osaka, Japan; # 010-01341) solution was prepared using PBS, administered intraperitoneally at 60 μL/kg, and sacrificed after 48 h. In the liver resection experiment, a 2/3 hepatic partial hepatectomy was performed using a previously reported protocol^37^. All studies were carried out following the Guide for the Care and Use of Laboratory Animals (National Institutes of Health) with the approval of Kyushu University’s Animal Care Committee.

### Human samples

The patients had previous admissions to Kyushu University Hospital for acute hepatitis, and histological examination revealed liver damage with necrosis. This study followed the Declaration of Helsinki and was approved by the Ethics Committee of Kyushu University Hospital (No. 23202-01). This was a retrospective study, so written informed consent was waived.

### Blood sample and biochemical analysis

Serum alanine aminotransferase (ALT) levels were determined using the DRI-CHEM NS500sV (FUJIFILM, Tokyo, Japan).

### Histological analysis

The livers of mice were fixed in 10% formalin for paraffin embedding and 4% paraformaldehyde for O.C.T compound embedding. Before hematoxylin and eosin (HE) staining, the livers were embedded in paraffin. To perform immunohistostaining, the samples were embedded in O.C.T. compound and serially cut into 10-μm sections. Human livers were fixed with 10% formalin and paraffin-embedded. Following antigen retrieval, the sections were blocked and incubated with primary antibodies. The primary antibodies used were anti-c-jun (Cell Signaling Technology; # 9165), anti-TP53inp1 (Abcam; # ab202026), anti-Ki67 (Invitrogen; # 14-5698-82), anti-cyclinD1 (Abcam; # ab134175), and anti-cyclinB1 (Cell Signaling Technology; # 4138). The primary antibodies were diluted from 1:100 to 1:200. The immunofluorescence staining was performed using Alexa Fluor 594- and 488-conjugated secondary antibodies (Invitrogen; # A11008, # A11006, # A11012, # A11007) at a dilution of 1:300. Immunohistochemistry was performed with N-Histofine Simple Stain Mouse MAX PO R (Nichirei Biosciences Inc, Tokyo, Japan; # 414341). Phalloidin-iFluor 594 Conjugate (Cayman; # 20553) was diluted at 1:1000. Finally, the sections were examined using a Keyence BZ-X700 microscope (Keyence, Osaka, Japan). Image analysis was done using ImageJ software (National Institutes of Health, Bethesda, MD). The DAPI integrated density was corrected by dividing the RawIntDen of the nucleus by the average luminance of three pixels surrounding the extracted nucleus. The corrected DAPI integrated density was visualized with the ggplot2 package (3.4.4) in R (4.3.2).

### Quantitative reverse transcription polymerase chain reaction

The total RNA from liver tissue was extracted with ISOGEN II reagent (NIPPON GENE CO., Ltd., Tokyo, Japan; # 311-07361), and cDNA was synthesized with the PrimeScript RT Master Mix kit (Takara Bio, Tokyo, Japan; # RR036A). The quantitative reverse transcription polymerase chain reactions (RT-qPCRs) were conducted using the THUNDERBIRD SYBR qPCR Mix (Toyobo Co., Ltd., Osaka, Japan; # QPX-201). The target gene expression was normalized to glyceraldehyde 3-phosphate dehydrogenase (*Gapdh*) expression, and relative expression was determined using the 2−ΔCt method. The primer sequences used in this study are *Jun*; Forward (5’–3’): GTGCCTACGGCTACAGTAAC, Reverse (5’–3’): CGACGTGAGAAGGTCCGAGTT, *Egfr; Forward* (5’–3’): AGTGGTCCTTGGGAACTTGG, Reverse (5’–3’): TGAGGGCAATGAGGACATAGC, and *Gapdh*; Forward (5’–3’): TGTGTCCGTCGTGGATCTGA, Reverse (5’–3’): TTGCTGTTGAAGTCGCAGGAG.

### Western blotting

The liver tissues were homogenized using RIPA buffer (NACALAI TESQUE, INC., Kyoto, Japan; # 16488-34). Total protein (30 μg) from each sample was loaded onto a Mini-PROTEAN TGX Gels (Bio-Rad Laboratories) (EGFR:4-15% (# 4561086), c-jun:7.5% (# 4561026)). Following electrophoresis, the proteins were transferred to a polyvinylidene difluoride membrane (Bio-Rad Laboratories, # 1704156) using the Trans-Blot Turbo Transfer System (Bio-Rad Laboratories). The membrane was incubated overnight at 4°C using a primary antibody. The primary antibodies used were anti-phospho-c-jun (Cell Signaling Technology; # 3270), anti-c-jun (Cell Signaling Technology; # 9165), anti-EGFR (Abcam; # ab52894), anti-phospho EGFR (Cell Signaling Technology; # 3777), and anti-GAPDH (Cell Signaling Technology; # 2118S). The primary antibodies were diluted at 1:1,000 dilution except for GAPDH, which was diluted at 1:2000 dilution. Then, the membrane was incubated with goat anti-rabbit IgG HRP-conjugated polyclonal antibody (Cell Signaling Technology; #7074S) at room temperature. The antibody was used in a 1:1,000 dilution. Proteins were detected using the Western Lightning ECL Pro (Revvity Japan Co., Ltd., Kanagawa, Japan; # NEL120001EA) and visualized with a ChemiDoc Imaging System (Bio-Rad Laboratories). Signals were quantified using ImageJ software (National Institutes of Health, Bethesda, MD, USA).

### Single-cell RNA sequencing analysis

scRNA-seq of APAP-treated mice was carried out using previously published data from Shani Ben-Moshe et al. ^7^. Hepatocyte data from m1-48 h and m1-72 h were analyzed. In addition to Shani Ben-Moshe et al., hepatocyte data (ZT6A and ZT6B) published by Droin et al. were used for 0 h samples ^38^. All analyses were carried out using the Seurat package (4.4.0) in R (4.3.2). Following normalization with NormalizeData (normalization. method = "LogNormalize," scale. factor = 10,000), each sample was integrated using genes identified by FindIntegrationAnchors. Doublets and low-quality cells were excluded by setting a total UMI count threshold of 1000-3500 and a mitochondrial gene threshold of less than 20%. They were then clustered using Seurat’s functions FindNeighbors (dims = 1:20) and FindClusters (resolution = 1.2). The cell cycle was determined using CellCycleScoring. AverageExpression was used to determine the average gene expression values. Signal scores were calculated with AddModuleScore. Gene ontology enrichment analysis was carried out using the functions enrichGO and simplify of the clusterProfiler package (4.10.0), with terms corresponding to BPs. To analyze stemness, a heat map was created using genes from KEGG’s signal pathways regulating pluripotency of stem cells (mmu04550) that are expressed in more than 10% of cells in some clusters.

The scRNA-seq of partially hepatectomized mice was carried out using publicly available data from Chembazhi. et al. ^6^. Following normalization with NormalizeData (normalization. Method = "LogNormalize," scale. factor = 10,000), each sample was integrated using genes identified with FindIntegrationAnchors. Doublet and low-quality cells were filtered using thresholds of <2,500 total UMI count, <2000 expressed gene count, and <10% mitochondrial gene. Because the total number of cells was 247,483 cells (40000-60000 cells each time), 5,000 cells from each time were randomly selected for analysis. Clustering was conducted using FindNeighbors (dims = 1:15) and FindClusters (resolution = 1.2). The cluster7 score was calculated by AddModuleScore which included 84 of the cluster7 and 13 characteristic genes found using FindMarkers that were upregulated in cluster7 and 13 and had avg_log2FC>1. The same analysis was repeated three times to ensure reproducibility.

### Statistical analysis

The data were analyzed with GraphPad Prism (version 10.1.1) (GraphPad Software, Inc.). The results were presented as means and standard deviations. The Mann-Whitney test was used to see if the difference between the two groups was significant. When comparing means across multiple groups, one-way ANOVA analysis and the Tukey test were used to identify significant differences.

P-values below 0.05 were considered statistically significant.

---

**Supplemental Figure 1.**
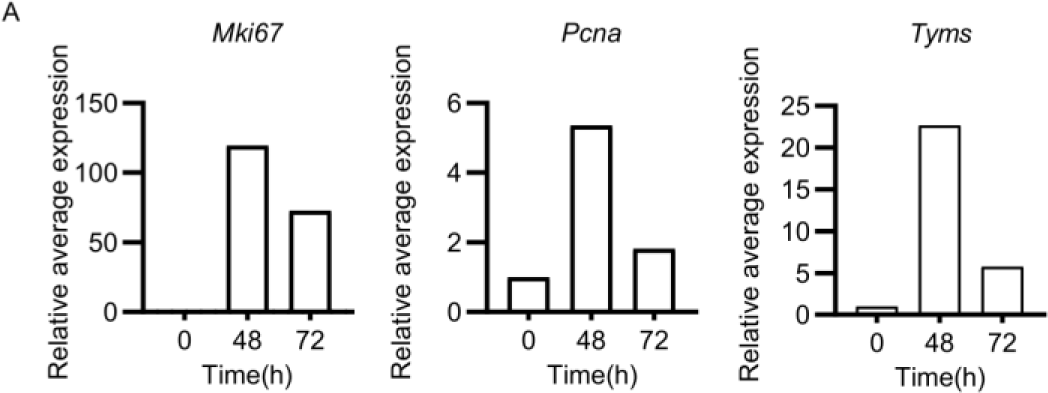
(A) The relative value of the average expression value for each hour compared to 0 h.

**Supplemental Figure 2.**
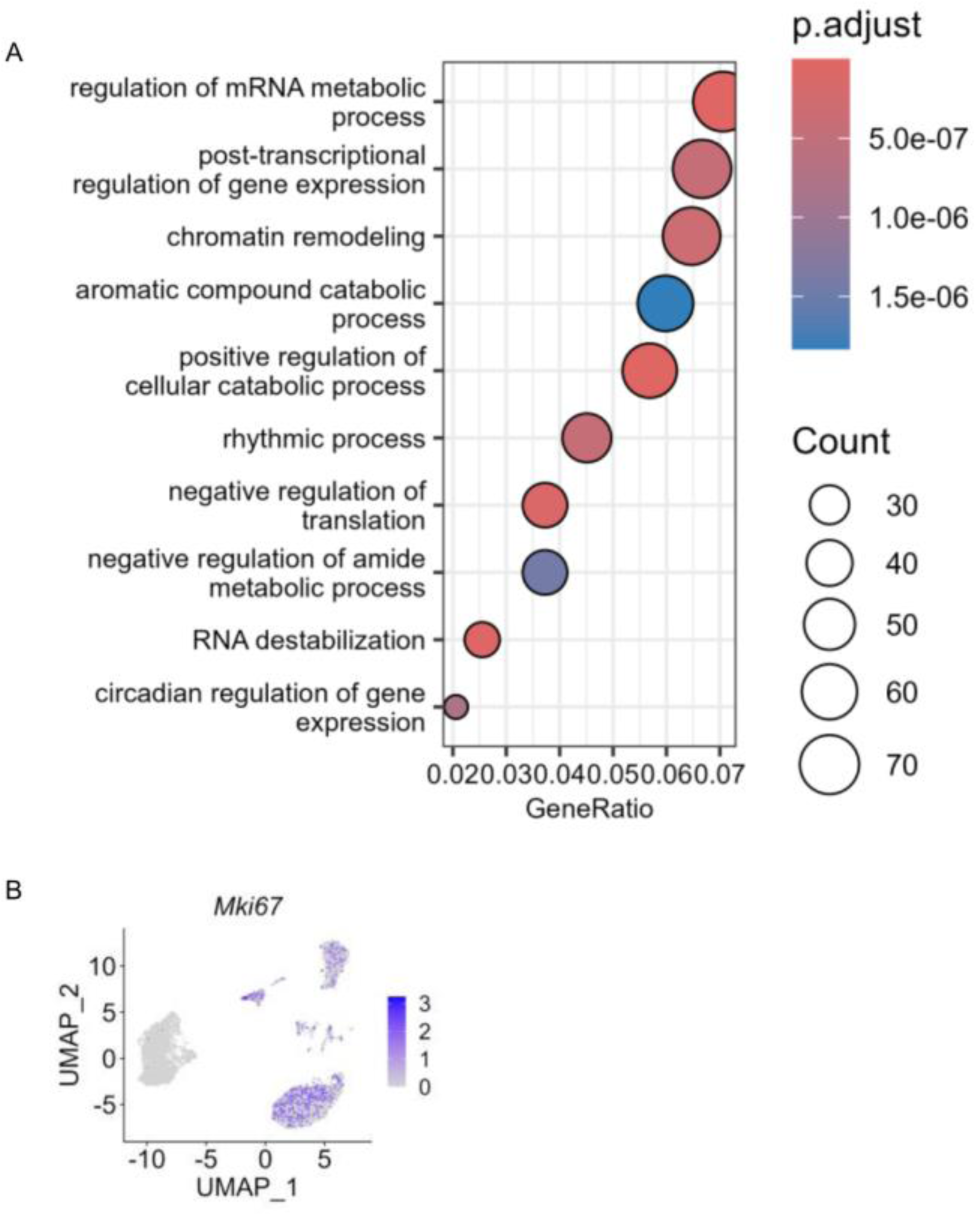
(A) Gene ontology enrichment analysis of genes that are up-regulated in cluster 7. (B) UMAP depicting the expression levels of Mki67

**Supplemental Figure 3.**
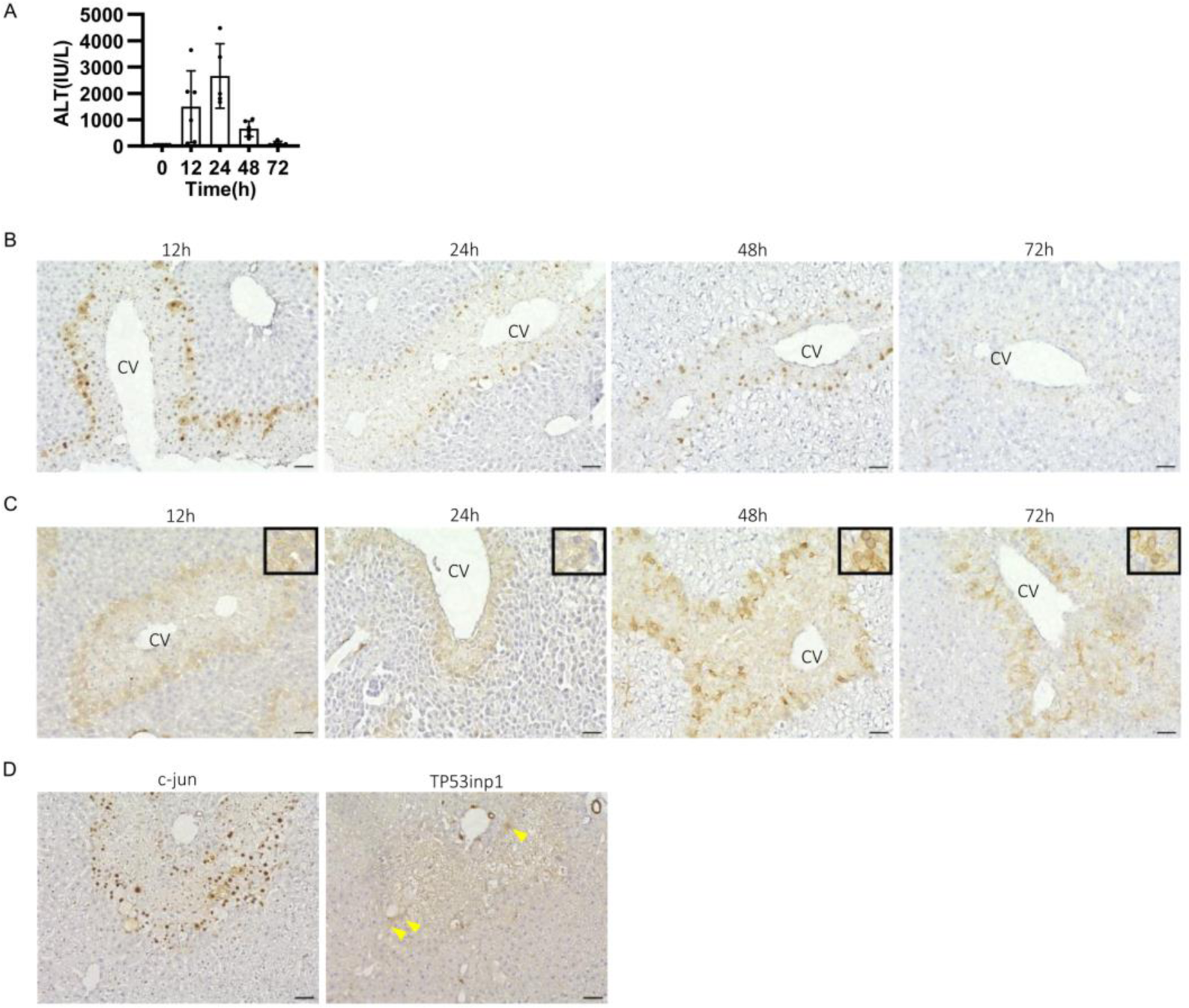
(A) The ALT for each time point is shown. (B) Immunohistochemical staining of c-jun. Scale bar is 50 μm. CV refers to the Central Vein. (C) Immunohistochemical staining for TP53inp1. The inset shows a magnified image of representative positive cells. Scale bar is 50 μm. (D) Immunohistochemical staining of c-jun and TP53inp1 in mice treated with allyl alcohol (sacrificed after 48h). The arrowheads indicate representative positive cells. Scale bar is 50 μm. PV refers to the Portal Vein.

**Supplemental Figure 4.**
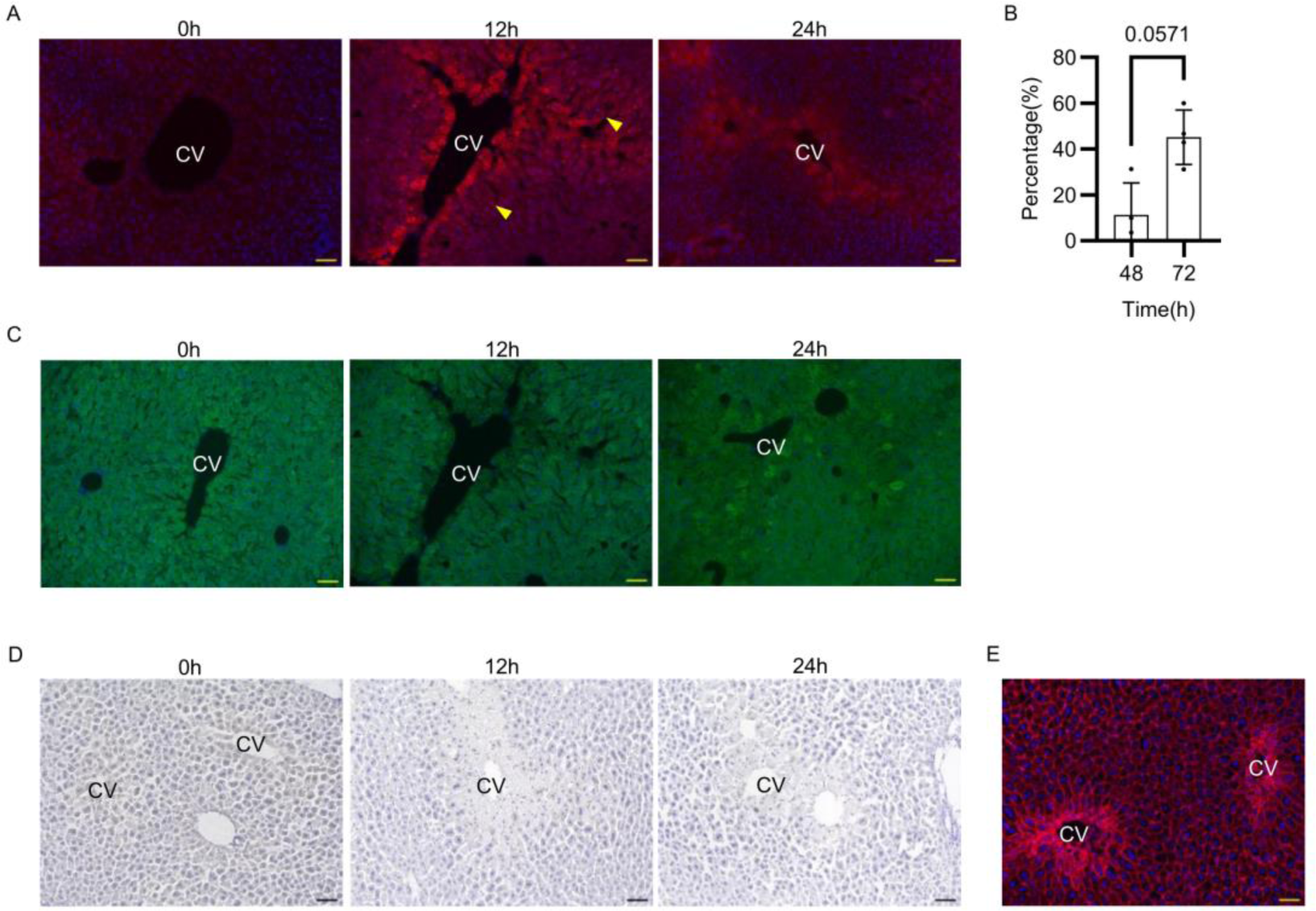
(A) Immunofluorescence staining of Ki67 from 0h to 24 h. The arrowheads indicate representative positive cells. TxRed, Ki67. Blue,DAPI. Scale bar: 50 μm. (B) Ki67 positivity in TP53inp1-positive cells. (C) Immunofluorescence staining of cyclinD1 from 0h to 24 h. Scale bar is 50 μm. (D) Immunohistochemical staining of cyclinB1 from 0 h to 24 h. The scale bar is 50μm. (E) A representative fluorescence staining of phalloidin. The sample is from 48h. TxRed,Phalloidin. Blue,DAPI. The scale bar is 50 μm

**Supplemental Figure 5.**
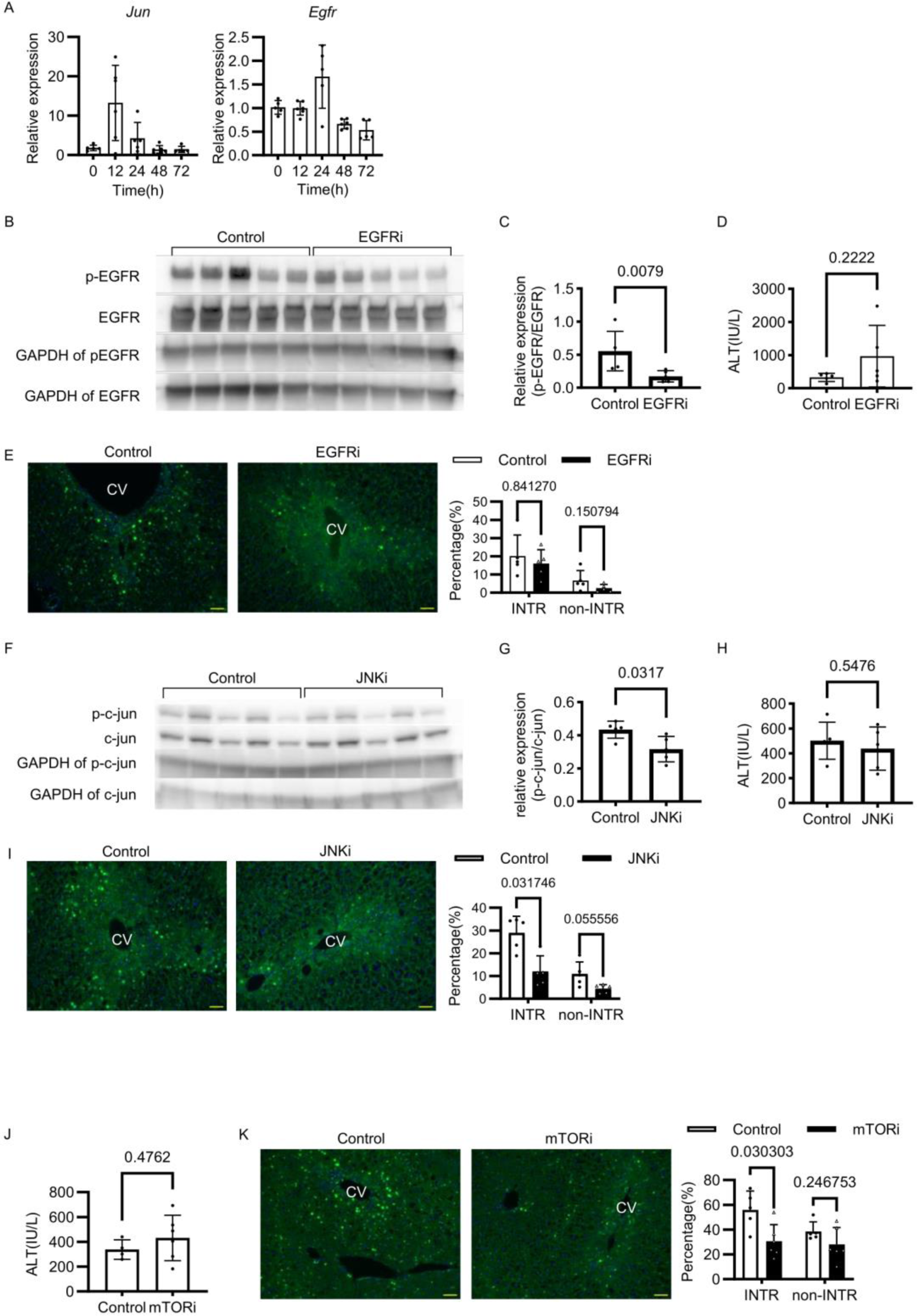
(A) The relative mRNA expressions of *Jun* and *Egfr* are shown at each time point. Normalized by *Gapdh*. (B) Western blots of phosphorylated EGFR and EGFR from the Control and EGFRi groups. (C) The ratio of signal values for phosphorylated EGFR and EGFR in Western blots, normalized by GAPDH. (D) Serum ALT for the Control and EGFRi groups. (E) Immunofluorescence staining of cyclinD1 in EGFR inhibition experiment. GFP, cyclinD1. Blue, DAPI. The scale bar is 50 μm. The bars represent the percentage of cyclinD1 positivity. (F) Western blot analysis of phosphorylated c-jun and c-jun in the Control and JNKi groups. (G) The ratio of signal values for phosphorylated c-jun and c-jun in Western blots, normalized by GAPDH. (H) Serum ALT for the Control and JNKi groups. (I) Immunofluorescence staining of cyclinD1 in JNK inhibition experiment. GFP, cyclinD1. Blue, DAPI. The scale bar is 50 μm. The bars represent the percentage of cyclinD1 positivity. (J) Serum ALT for the Control and mTORi groups. (K) Immunofluorescence staining of cyclinD1 in mTOR inhibition experiment. GFP, cyclinD1. Blue, DAPI. The scale bar is 50 μm. The bars represent the percentage of cyclinD1 positivity.

**Supplemental Figure 6.**
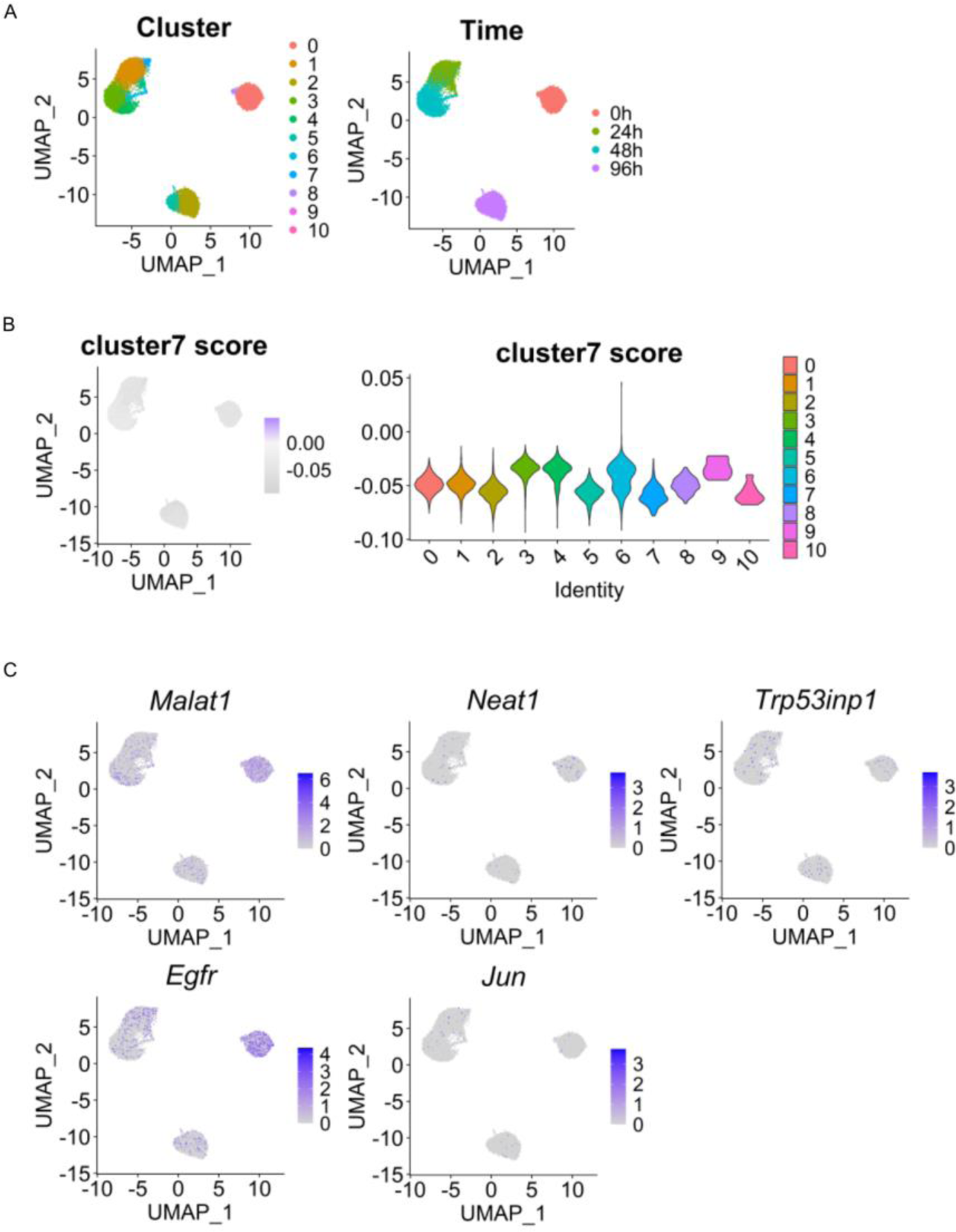
(A) UMAP of scRNA-seq in partially hepatectomized mice. (B) UMAP and violin plots with cluster7 score. (C)UMAP depicts the expression levels of each gene.

