## Supplemental Figure for "Two types of regeneration mechanism in acute liver injury"

A

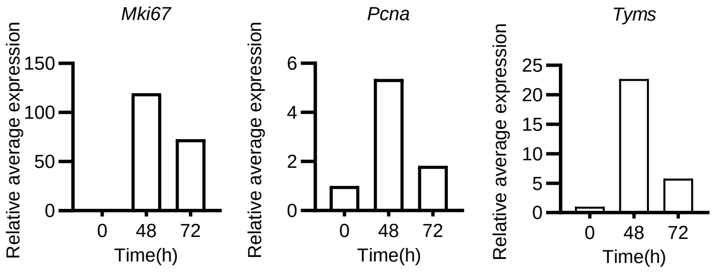


***Supplemental Figure 1***

(A)The relative value of the average expression value for each hour compared to 0 h.

A

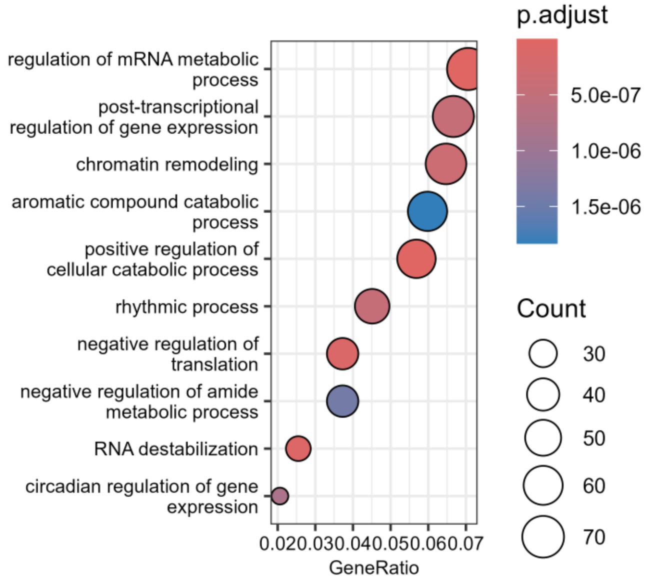


B

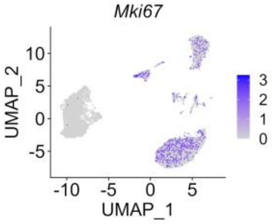


***Supplemental Figure 2***

(A) Gene ontology enrichment analysis of genes that are up-regulated in cluster 7. (B) UMAP depicting the expression levels of Mki67

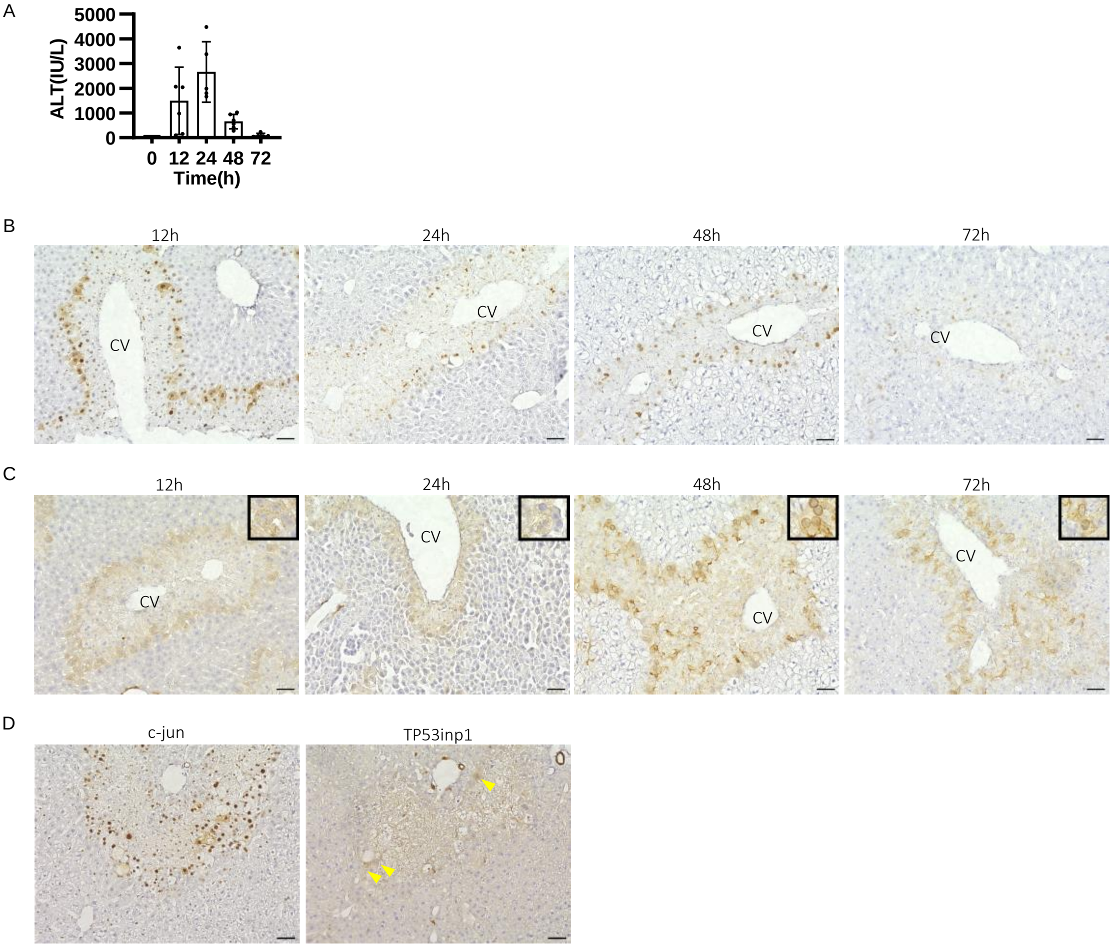


***Supplemental Figure 3***

(A) The ALT for each time point is shown. (B) Immunohistochemical staining of c-jun. Scale bar is 50 μm. CV refers to the Central Vein. (C) Immunohistochemical staining for TP53inp1. The inset shows a magnified image of representative positive cells. Scale bar is 50 μm. (D) Immunohistochemical staining of c-jun and TP53inp1 in mice treated with allyl alcohol (sacrificed after 48h). The arrowheads indicate representative positive cells. Scale bar is 50 μm. PV refers to the Portal Vein.

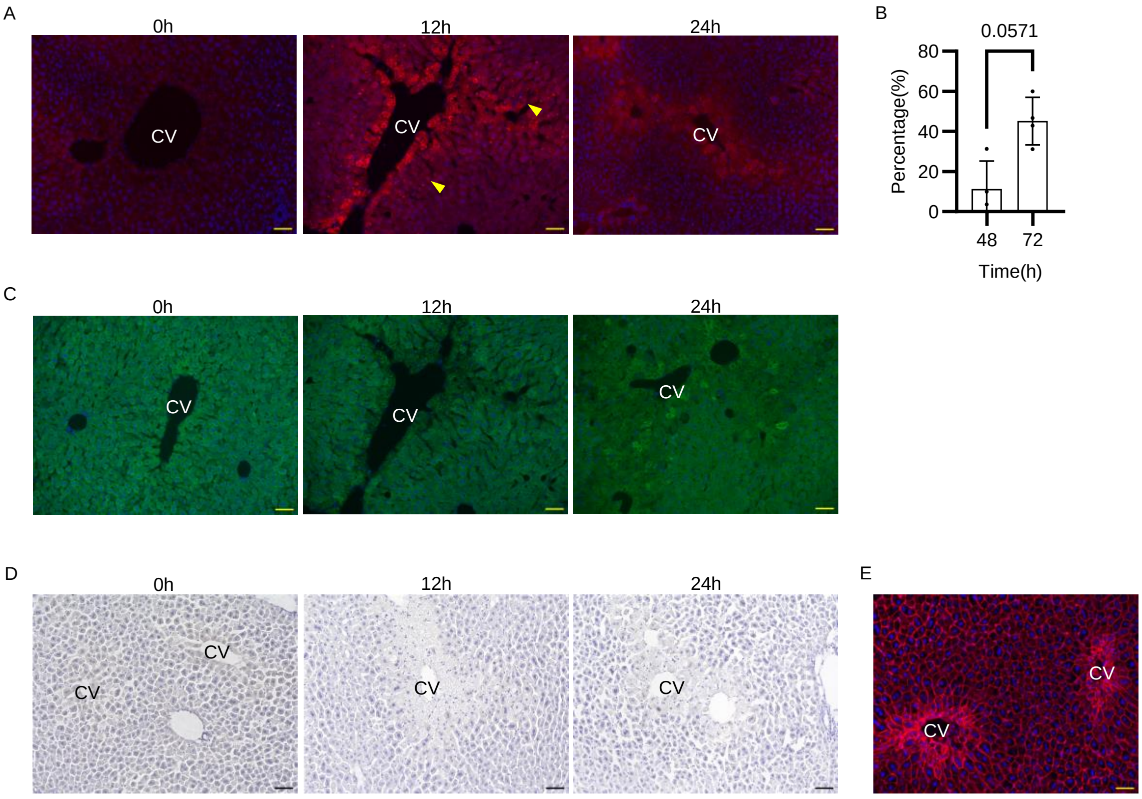


***Supplemental Figure 4***

(A) Immunofluorescence staining of Ki67 from 0h to 24 h. The arrowheads indicate representative positive cells. TxRed, Ki67. Blue,DAPI. Scale bar: 50 μm. (B) Ki67 positivity in TP53inp1-positive cells. (C) Immunofluorescence staining of cyclinD1 from 0h to 24 h. Scale bar is 50 μm. (D) Immunohistochemical staining of cyclinB1 from 0 h to 24 h. The scale bar is 50μm. (E) A representative fluorescence staining of phalloidin. The sample is from 48h. TxRed,Phalloidin. Blue,DAPI. The scale bar is 50 μm

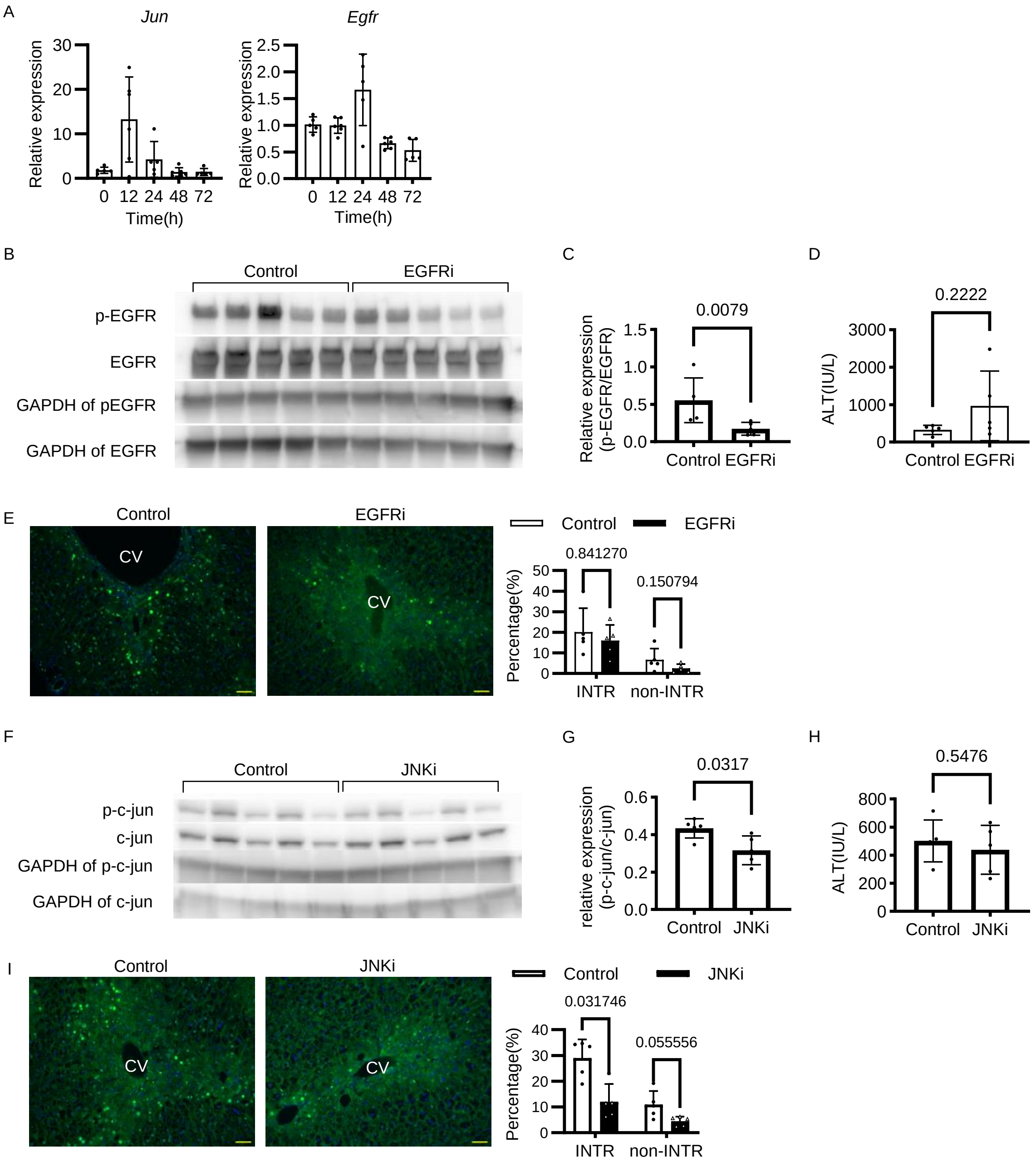


***Supplemental Fig.5***

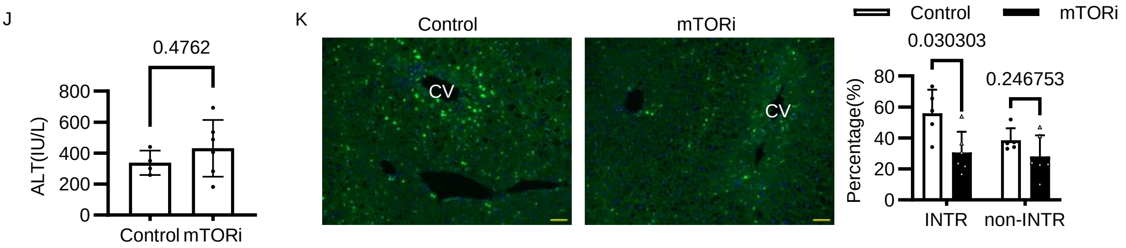


***Supplemental Figure 5***

(A) The relative mRNA expressions of Jun and Egfr are shown at each time point. Normalized by Gapdh. (B) Western blots of phosphorylated EGFR and EGFR from the Control and EGFRi groups. (C) The ratio of signal values for phosphorylated EGFR and EGFR in Western blots, normalized by GAPDH. (D) Serum ALT for the Control and EGFRi groups. (E) Immunofluorescence staining of cyclinD1 in EGFR inhibition experiment. GFP, cyclinD1. Blue, DAPI. The scale bar is 50 μm. The bars represent the percentage of cyclinD1 positivity. (F) Western blot analysis of phosphorylated c-jun and c-jun in the Control and JNKi groups. (G) The ratio of signal values for phosphorylated c-jun and c-jun in Western blots, normalized by GAPDH. (H) Serum ALT for the Control and JNKi groups. (I) Immunofluorescence staining of cyclinD1 in JNK inhibition experiment. GFP, cyclinD1. Blue, DAPI. The scale bar is 50 μm. The bars represent the percentage of cyclinD1 positivity. (J) Serum ALT for the Control and mTORi groups. (K) Immunofluorescence staining of cyclinD1 in mTOR inhibition experiment. GFP, cyclinD1. Blue, DAPI. The scale bar is 50 μm. The bars represent the percentage of cyclinD1 positivity.

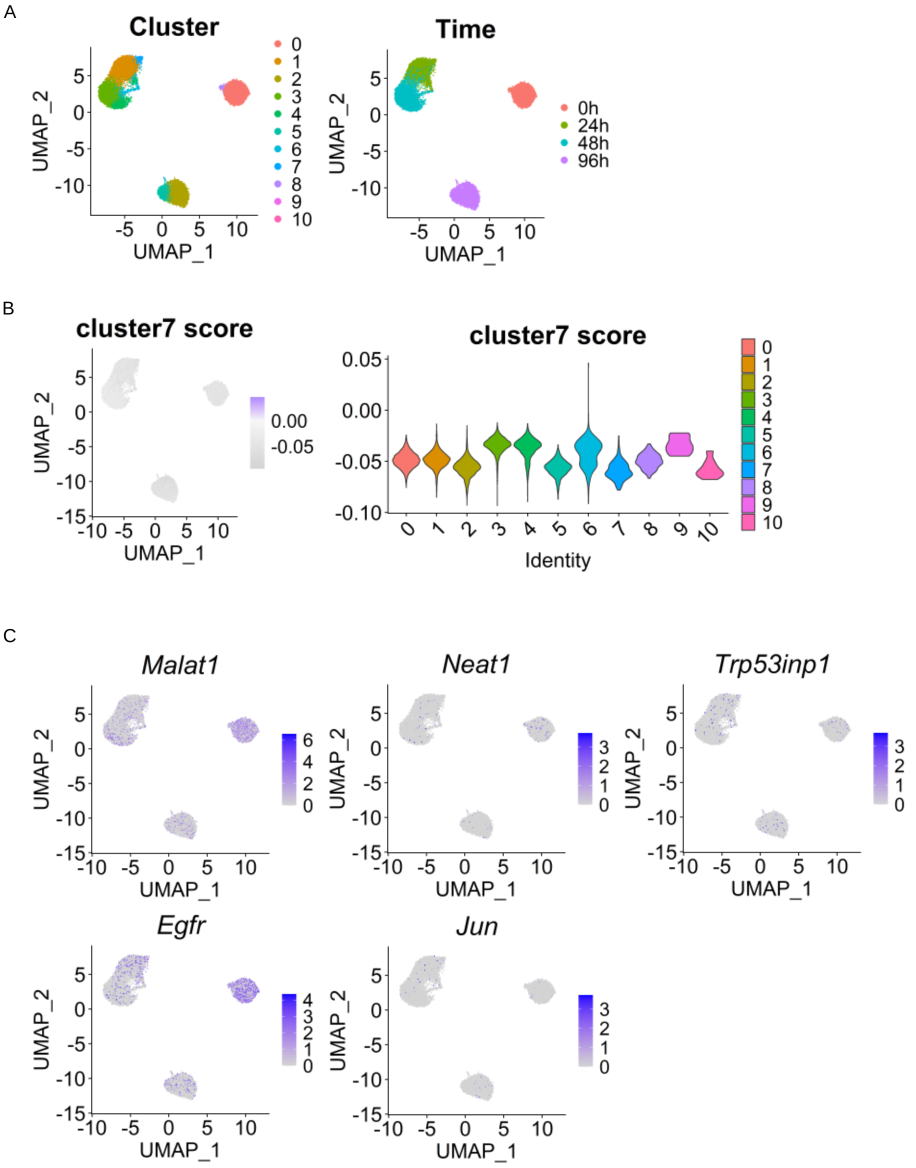


***Supplemental Figure 6***

(A) UMAP of scRNA-seq in partially hepatectomized mice. (B) UMAP and violin plots with cluster7 score.

(C)UMAP depicts the expression levels of each gene.
